# A phage display approach to identify highly selective covalent binders

**DOI:** 10.1101/791533

**Authors:** Shiyu Chen, Matthew Bogyo

**Affiliations:** Department of Pathology, Stanford University School of Medicine, Stanford, CA 94305, USA; Department of Microbiology and Immunology, Stanford University School of Medicine, Stanford, CA 94305, USA

**Keywords:** Phage display, covalent binders, activity-based probe, protease

## Abstract

Molecules that bind macromolecular targets through direct covalent modification have found widespread applications as activity-based probes (ABPs) and as irreversible drugs. Covalent binders can be used to dynamically monitor the activity of enzymes in complex cellular environments, identify targets and induce permanent binding/inhibition of therapeutically important biomolecules. However, the general reactivity of the electrophiles needed for covalent bond formation makes control of selectivity difficult. There is currently no rapid, robust and unbiased screening method to identify new classes of covalent binding ligands from highly diverse pools of candidate molecules. Here we describe the development of a phage display method to screen for highly selective covalent binding ligands. This approach makes use of a reactive linker to form cyclic peptides on the phage surface while simultaneously introducing an electrophilic ‘warhead’ to covalently react with a nucleophile on the target. Using this approach, we identified cyclic peptides that selectively and irreversibly inhibited a cysteine protease with nanomolar potency, exceptional specificity and increased serum stability compared to a linear peptide containing the same electrophile. This approach should enable rapid, unbiased screening to identify new classes of highly selective covalent binding ligands for diverse molecular targets.

## Introduction

A large portion of therapeutic agents mediate their activity through interactions with specific biomolecules. Efficacy can often be linked to the ability of the molecule to bind a single target with high selectivity. While drug discovery efforts have historically focused on reversible binding molecules, recent success with covalent modifiers has highlighted their great potential to be used as therapeutic agents[1, 2]. In addition, covalent modifiers of the active site residues of enzymes have found widespread use as activity-based probes (ABPs) for functionally characterizing and identifying diverse families of enzymes within native cellular environments[3, 4]. However, the use of chemically reactive electrophiles to mediate covalent binding interactions with nucleophilic residues (i.e. cysteine, serine and lysine) on proteins introduces the significant challenge of controlling selectivity. This potential liability of covalent binding molecules can be overcome by using weakly reactive electrophiles linked to high affinity binding elements that limit off-target interactions. However, the identification of binding components with sufficiently selectivity to pair with a reactive electrophile often requires painstaking efforts that are time consuming and potentially of limited success.

Over the past two decades, covalent binding ABPs that target a number of diverse classes of enzymes including proteases[3, 5, 6], lipases[7], esterases[8], kinases[9, 10], phosphatases[11] and glycosidases[12] have been reported. While a small number of ABPs have been reported that target a single enzyme[13-16], in most cases, they react with a sub-set of related enzymes[17, 18]. This is largely due to the highly conserved active site regions within many enzyme family members and the use of small molecule or peptide-like targeting sequences with limited surface area contacts between probe and target. Thus, there is a need to develop a general approach that will allow direct screening of large pools of highly structurally diverse covalent probes to identify selective agents for a given target of interest. Phage display is an ideal technology for this application as it allows rapid generation of billions of peptides of exceptionally high diversity on the surface of phage particles, which can then be iteratively screened against molecular targets to identify highly selective binders[19, 20]. The limitation of classical phage methods is that only simple linear peptides made up of natural amino acid sequence can be screened and the resulting hits are reversible binding ligands.

We describe here a general, unbiased approach to directly screen pools of covalent probes with exceptional diversity using the power of phage display. Inspired by the work of others in which phage are chemically modified to produce cyclic and bicyclic peptides directly on the phage surface[21-24], we developed a small molecule linker that could be used to both form cyclic peptides and also introduce a weak electrophile ‘warhead’ directly on the cyclized phage coat protein segment (Fig. 1). This allowed iterative panning of phage containing large pools of potential covalent binding cyclic peptides. Cyclization helps to both rigidify the peptide scaffold for increased binding potency and selectivity and also to increase the overall biological stability of the resulting probes. These types of cyclized peptides also tend to adopt secondary structures that more closely mimic a folded protein and therefore are capable of binding with an increased surface area of contacts compared to a linear peptide[25]. This greatly increases the chances of identifying ligands with highly selective binding properties. We chose to validate the approach using Tobacco Etch Virus (TEV) protease since it is known to have well-defined primary sequence selectivity yet there are currently no reported potent inhibitors of this target[26]. Our screening efforts using a cyclization linker containing a vinyl sulfone warhead identified a series of peptide sequences that irreversibly inhibited TEV only when cyclized. The consensus sequences of phage selected peptides contained residues found in the P3-P6 residues of the native TEV substrate. However, the critical P1 and P2 residues required for efficient targeting of TEV were missing, indicating an alternate mode of binding compared to a native substrate. Molecular modeling confirmed an energy minimized structure in which the cyclic peptide portion of the molecule displays the key P3, P4 and P6 residues of the native substrate when the reactive vinyl sulfone is covalently linked to the active site cysteine residue. Further optimization of the primary sequence of the peptide scaffold from phage panning resulted in a selective covalent inhibitor of TEV with nanomolar potency. Attachment of a fluorescent dye produced an ABP for TEV that specifically labeled the enzyme in complex proteomic mixtures. This probe and parent cyclic peptide -VS inhibitor showed dramatically increased target selectivity as well as serum stability compared to a corresponding linear peptide vinyl sulfone inhibitor based on the natural substrate recognition sequence for TEV. Therefore, we were able to use phage display to identify rigid cyclic peptide structures that presented an attached electrophile to the active site cysteine of a protease to yield a covalent inhibitor and ABP with exceptional selectivity. This phage screening approach has the potential to be a general method for rapid screening to identify highly selective and stable covalent binding molecules for any molecular target that contains a suitable nucleophile.

**Fig. 1.**
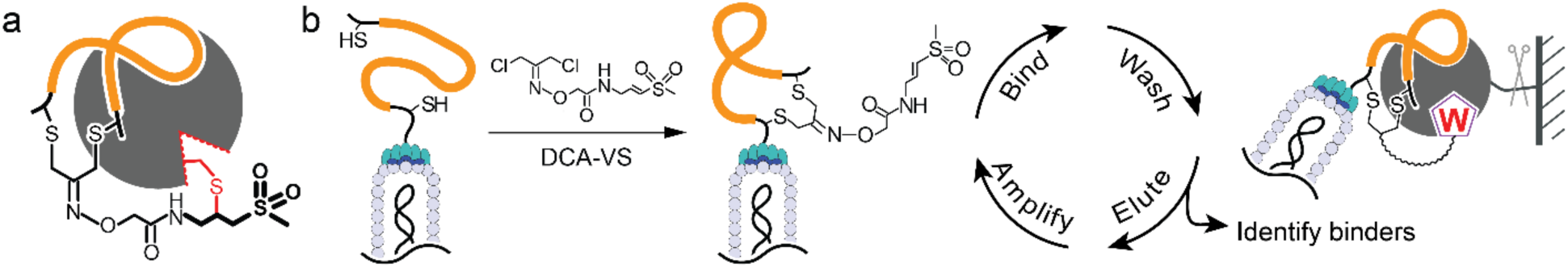
The Phage display approach to screen for selective covalently binding ligands. a) Cartoon of the general strategy in which a linker containing the reactive vinyl sulfone (black) is used to cyclize peptides (orange) such that they can be presented to a target by binding in the proximity of a reactive nucleophile (red). b) General workflow of the phage screening method to find selective covalent binding ligands. The DCA-VS linker is used to chemically modify diverse peptides containing two cysteines to generate a cyclic peptide-vinyl sulfone library on the phage surface. The resulting library can be screened through an iterative process in which phage that covalently bind the target are collected by affinity purification followed by washing to remove non-covalently bound phage and then elution by proteolytic digestion of the target linker. The resulting released phage can then be amplified and subjected to the next round screening.

## Results

### Design of the cyclization linker

We set out to develop a linker that could be used to both form constrained cyclic peptides while simultaneously introducing a reactive electrophile to produce potential covalent binding probes directly on the phage surface. This general approach was inspired by the pioneering work by Heinis and Derda using chemical cross-linker to generate phage-displayed monocyclic/bicyclic peptides to screen for highly selective inhibitors of protease targets[21, 23]. This work showed that by constraining the peptide as a bicyclic peptide, it was possible to generate diverse ‘protein-like’ structures that formed large surface area contacts with the target protein, resulting in inhibitors with exceptional target selectivity and potency. In order to identify such constrained structures of high affinity and selectivity it is necessary to perform selections with pools of phage with highly diverse peptide sequences. We therefore focused on identifying a linker that could efficiently induce cyclization while carrying a weakly reactive electrophile.

1,3-Dichloroacetone (DCA) has been reported to be a highly selective linker for cyclizing peptides in solution through reaction with two cysteines, as well as directly modifying phage displayed peptides without causing toxicity[24, 27]. The ketone group of DCA allows convenient derivatization with alkoxy-amines/hydrazine to generate stable oxime/hydrazone linkages for introducing various functional groups without disrupting the α-halo groups needed for cyclizing peptides[28]. Cyclization through reaction with two free thiols on a peptide occurs rapidly and selectively under physiologically relevant conditions (pH 6.5-7.5). We synthesized a suitable linker by coupling tert-Butyloxycarbonyl (Boc) protected aminooxy acetate to the N-terminus of glycine-vinyl sulfone followed by deprotection in trifluoroacetic acid (TFA) to produce the free aminooxy group which then quantitatively reacted with DCA at room temperature to produce the desired DCA-VS linker (**11**, Supplement Scheme 1) in good yield.

Among the various electrophilic warheads that have been used in ABPs to target cysteine proteases, we chose the vinyl sulfone due to its high level of selectivity for thiol nucleophiles[29]. Furthermore, the α-amino vinyl sulfone warhead has overall low reactivity towards free cysteine and can only target the highly nucleophilic active site cysteine of cysteine proteases in the presence of a guiding peptide sequence to orientate the warhead in the active site. We also chose to use a simple glycine vinyl sulfone (**11**, Supplement Scheme 1) because we did not want selection to be driven by the P1 residue containing the electrophile but rather by the cyclic peptide diversity portion of the molecule. We reasoned that rigid, protein like cyclic peptide structures could be identified that would position the warhead in proximity of the active site nucleophile and induce formation of the covalent bond (Fig. 1a). The general screening workflow using the DCA-VS linker involves treating a library of phage expressing diverse peptides containing two cysteines on their pIII coat to induce cyclic peptide and attach the VS electrophile. The resulting chemically modified phage can then be screened for covalent binding to a suitably labeled target protein with elution of hits by proteolytic cleavage from a solid support (Fig. 1b).

To first confirm that the DCA-VS linker forms the desired cyclization product on the phage coat protein, we applied the linker to cyclize a di-cysteine peptide CysX_8_Cys (^N^ACGSGSGSGCG^C^) fused to the N1 and N2 ectodomains of the cysteine-free phage pIII protein[30]. After reducing the cysteine-rich peptide with TCEP, we incubated the protein with the DCA-VS linker at 30 °C for 2 hours and monitored the cyclization reaction by mass spectrometry as has been reported[31]. These studies confirmed quantitative and specific cyclization by the DCA-VS linker in a phage compatible buffer (Fig. 2a).

**Fig. 2.**
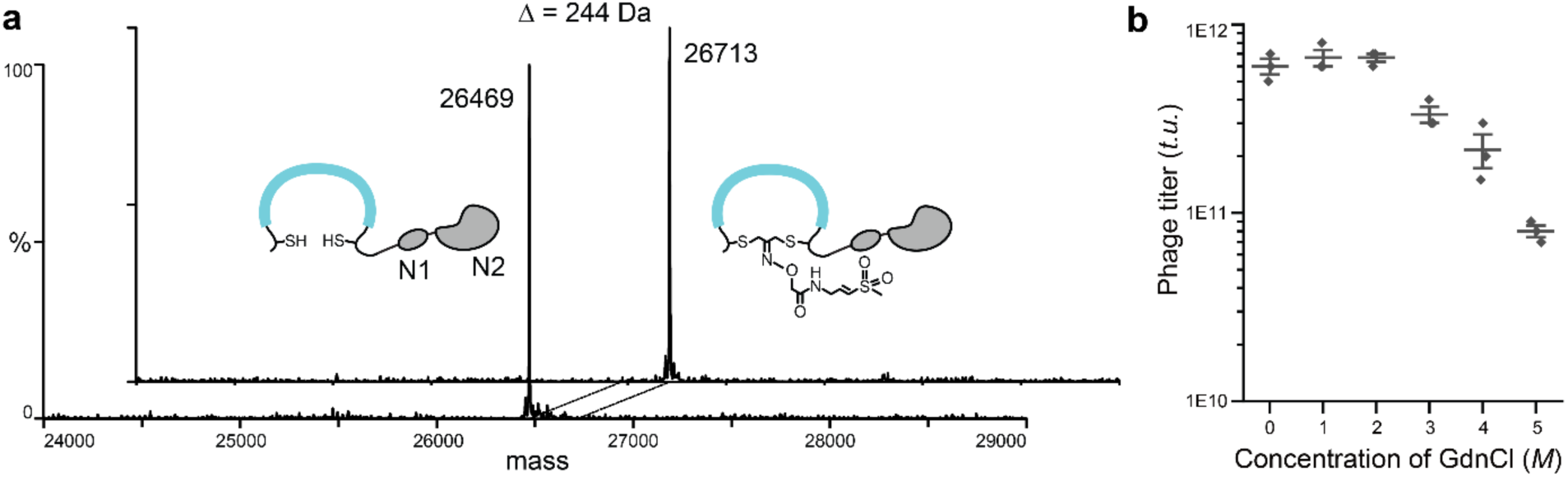
Confirmation of selectivity of the cyclization reaction and establishment of washing conditions for removal of non-covalently bound phage. a) A test peptide containing two cysteine residues (^N^A<u>C</u>GSGSGSG<u>C</u>G^C^) fused to the disulfide free phage N1N2 domains is quantitively modified to generate the desired cyclic peptide conjugate. The mass shift corresponds to the adduct of a single DCA-VS linker. b) Phage expressing a disulfide-free pIII protein were treated with increasing concentrations of guanidine chloride (GdnCl) to determine the highest concentration that could be used in washing steps without killing the phage. Assays were performed in triplicate, n= 3 biologically independent wells (mean + s.d. is depicted).

### Design of target protease construct and optimization of washing conditions

Because the linker introduces a reactive electrophile directly on the phage surface, the desired phage ‘hits’ will be covalently bound to the target protein. This provides an opportunity to remove reversibly bound as well as non-specifically bound phage using stringent washing conditions. Collection of the desired phage can then be achieved by selective release of the target from the resin using a protease cleavage site. Therefore, we needed to investigate washing conditions that would not harm the phage and also design a TEV protease construct that would allow release by a protease. We engineered an expression construct containing an AviTag biotinylation tag linked to the N-terminus of TEV protease through a linker recognized by the Human Rhinovirus (HRV) 3C Protease[32]. To improve the solubility and folding of the TEV protease, we also fused *E. coli* maltose-binding protein (MBP) through a TEV cleavable sequence (ENLYFQG) N-terminal to the AviTag[33] (Supplement Fig. 1). During expression, the TEV protease autocleaves the MBP moiety yielding a soluble TEV protease containing both a hexahistidine tag for affinity purification and the cleavable biotin tag for releasing covalently bound phage. To identify optimal wash conditions that remove non-covalently bound phage, we tested a range of concentrations of guanidine. We found that the infectivity of phage remains high even after treatment with 5M guanidine chloride (Fig. 2b) and therefore used these washing conditions to remove reversibly bound phage during each stage of the phage screening process.

### Identification of covalent binding cyclic peptide probes

For our phage panning experiments we used a library of phage displaying peptides of eight random amino acids flanked by two cysteines (Cys-(Xxx)8-Cys, Xxx represents any canonical amino acid; Fig. 3a) inserted at the N-terminus of the disulfide-free pIII protein[34]. The randomized DNA sequences encoding the peptide library were inserted into the phage DNA between the pelB signal peptide and pIII protein of the fdg3p0ss phage vector[30]. The phage peptide library was produced in TG1 cells by transforming host bacteria, at 30 °C for 16 hours. To overcome the low infectivity of the disulfide-free phage strain used in our screens, we used a large volume of 2YT rich media (1 L) for production of phage. The cysteine rich peptides present on the phage surface were reduced with TCEP and then modified with 300 μM of the DCA-VS linker to achieve the cyclic peptide-vinyl sulfone (cpVS) phage library. To select for covalent binding phage, we immobilized the biotinylated TEV protease on streptavidin magnetic beads and incubated at room temperature to allow a complete reaction between the active phage and the TEV protease. To promote stronger and faster binding upon increasing rounds of selection, we allowed phage and target to bind for 4 hours in the first round of selection, for one hour in the second round, and a half hour in the third round. Measurement of the overall phage titer levels indicated a slight reduction in phage numbers compared to conventional phage screening, likely due to the stringent 5M guanidine washes. However, we were able to achieve an approximately 55-fold enrichment in phage numbers through three rounds of panning with titers increasing from 1E4 to 5.5E5, suggesting that the selection was successful. To release the selected phage from solid support, we incubated the affinity resin with the 3C protease. Phage collected from the third round of selection were used to infect TG1 *E. coli*, that were then grown on plates to produce individual colonies for sequencing of recovered plasmid DNA.

**Fig. 3.**
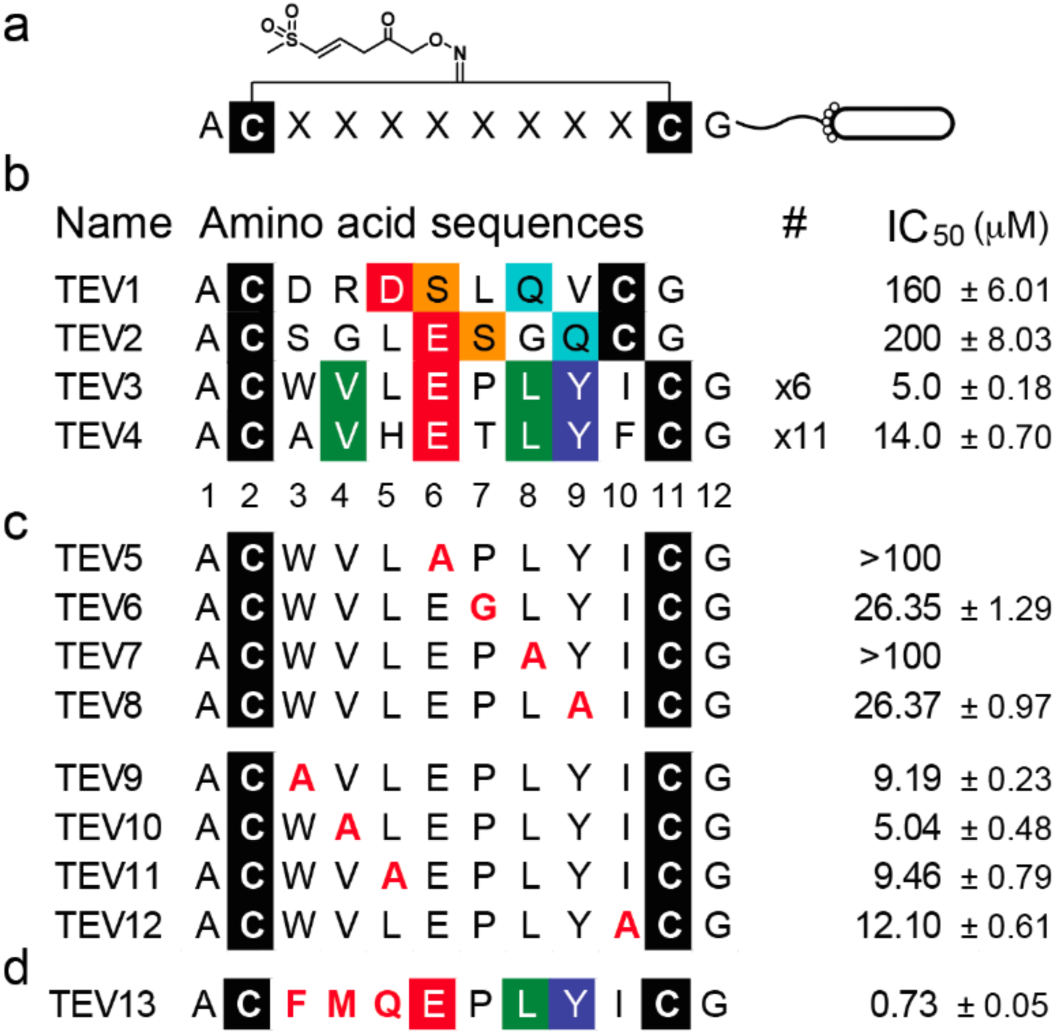
Identification and optimization of covalent cyclic peptide inhibitors of TEV protease. a) Cartoon of the cpVS phage library, generated by cyclizing linear peptides containing two cysteine residues separated by 8 diversity positions (Cys-(Xxx)8-Cys where Xxx represents any canonical amino acid) using the cysteine reactive DCA-VS linker. b) The primary sequences of cpVS inhibitors selected after three rounds of panning against the TEV protease. The identified sequences are grouped according to similarity in sequences. Conserved amino acids are colored using the Shapely coloring rules. For peptides found more than once, the frequency in a pool of 19 sequenced clones is indicated (#). Each sequence was chemically synthesized, converted into a cyclic peptide containing the DCA-VS linker and tested for potency against the recombinant TEV protease using a fluorogenic substrate assay. The numbering for the amino acid positions are shown (bottom). c) Sequences of peptides used for the mutational analysis to identify key binding residues. The top panel shows mutants in the conserved 6-9 positions. The lower panel shows mutations of the 3-5 and 10 position. The IC_50_ values for each peptide sequences when converted to the corresponding cpVS inhibitor are shown. d) Sequence and IC_50_ of the optimized cpVS TEV13 constructed based on the results from the mutational analysis in (c). The residues mutated are shown in red. The resulting average half maximal inhibitory concentrations (IC_50_) and standard deviations are shown, n= 3 biologically independent wells.

The sequencing results from 19 clones identified consensus peptide sequences that can be separated into two groups (Fig. 3b). The first group contained two peptides with a D/ESXQ sequence in the variable 7 residue stretch between the two cysteines (TEV1 and TEV2). The second group contained two peptides with a VXEXLY sequence in an 8 residue stretch between the two cysteines (TEV3 and TEV4). This sequence was isolated 17 times from the original 19 colonies. These results were encouraging because both sets of sequences contained conserved residues found in the established TEV protease substrate binding sequence EXLYXQ, yet both lack the key P1 and P2 residues that form contacts to the active site suggesting a likely alternate binding mode to a linear peptide sequence.

To validate the identified sequences from the phage screens, we synthesized all 4 consensus sequences as free peptides containing a free N-terminus and a C-terminal amide using standard solid phase peptide chemistry. We then produced the resulting cyclic peptides by reacting with the DCA-VS linker and purified the resulting cyclic peptide covalent inhibitors. To test the inhibitory potency of the cyclized peptides for TEV protease, we performed dose response analysis using a quenched fluorogenic peptide (Cy5-ENLYFQGK(QSY21)-NH_2_) as a reporter for protease activity. Peptides were preincubated with TEV for 1 hour at 30 °C followed by addition of substrate and measurement of Cy5 fluorescence increase over time. This allowed us to calculate an IC_50_ value for each peptide that could be used as a general metric for potency of each relative to one another. These results confirmed that the inhibitors containing the D/ESXQ sequence in a variable 7 residue stretch (TEV1 & TEV2) were an order of magnitude less potent that the inhibitors containing the VXEXLY sequence in an 8 residue stretch (TEV3 & TEV4). Importantly, the most potent inhibitor, TEV3, had an IC_50_ inhibition value of 5 μM, making it an ideal starting point for further optimization to achieve a high affinity TEV inhibitor.

### Optimization to generate a covalent TEV inhibitor with nanomolar potency

To determine which residues of the TEV3 sequence were most important for target binding, we replaced each residue at position 3-6 and 8-10 with alanine and used glycine to replace the proline at position 7 (Fig. 3c). After synthesizing each individual mutant peptide, cyclizing with DCA-VS (TEV5-TEV12) and testing activities against TEV using the substrate activity assay, we found that the conserved glutamic acid at position 6 and leucine at position 8 were essential for binding (TEV5 & TEV8). Mutation of the proline at position 7 and leucine at position 8 resulted in a dramatic drop in activity (TEV6 & TEV7). On the other hand, mutation of any of the other positions only slightly decreased inhibitor potency, suggesting that these positions can be altered to improve binding potency.

We then performed rational optimization by keeping the essential residues (position 6-9), while substituting the remaining residues in the peptides with natural amino acids of different sizes, hydrophobicity and electrostatic properties. As before, each peptide was synthesized on solid support, modified with the DCA-VS linker and purified to generate the corresponding cpVS conjugate (Supplement Fig. 2). For positions 3-5, phenylalanine at position 3, methionine at position 4 and glutamine at position 5 showed the most potent inhibition in each group (Supplement Fig. 2a-c). Modifications at position 10 did not improve potency so we fixed this position as the original isoleucine (Supplement Fig. S2d). Incorporation of the three preferred amino acids in positions 3-5 into a single peptide sequence resulted in the most potent inhibitor (TEV13) with nanomolar activity against the TEV protease (IC_50_ of 730 nM with 1-hour preincubation; Fig. 3d). This peptide was efficiently cyclized by the DCA-VS linker, showing no significant (<1%) free cysteine present in cyclic form of TEV13 (Supplement Fig. 3).

To demonstrate the selectivity of the optimized inhibitor, TEV13, we synthesized a number of control molecules. The first molecule is an isomer of TEV13 in which the amino acid residues between the two cysteines are scrambled to confirm the importance of the sequence order and rule out inhibition by generic reactivity of the VS electrophile (TEV14; Fig. 4a). The second control is the optimal TEV13 sequence that is cyclized with a linker that lacks the reactive VS electrophile (TEV15; Fig. 4a). This molecule allows assessment of the contribution of the cyclic peptide portion to binding in the absence of a covalent tether to the active site. As a final control we wanted to compare our optimal cyclic peptide covalent inhibitor to an inhibitor designed based on the defined substrate specificity of the target protease. TEV is ideal for this comparison since it is rather unique in that it has a highly defined extended sequence specificity resulting from multiple contacts between its substrate and the protease active site[35, 36]. Therefore, we synthesized a linear hepta-peptide vinyl sulfone containing the optimal TEV substrate sequence ENLYFQ (TEV16; Fig. 4a). This linear peptide vinyl sulfone probe was synthesized directly on solid support by linking the P1 amino acid vinyl sulfone through the glutamine side chain to the resin as described previously[37] (Supplement Scheme 2).

**Fig. 4.**
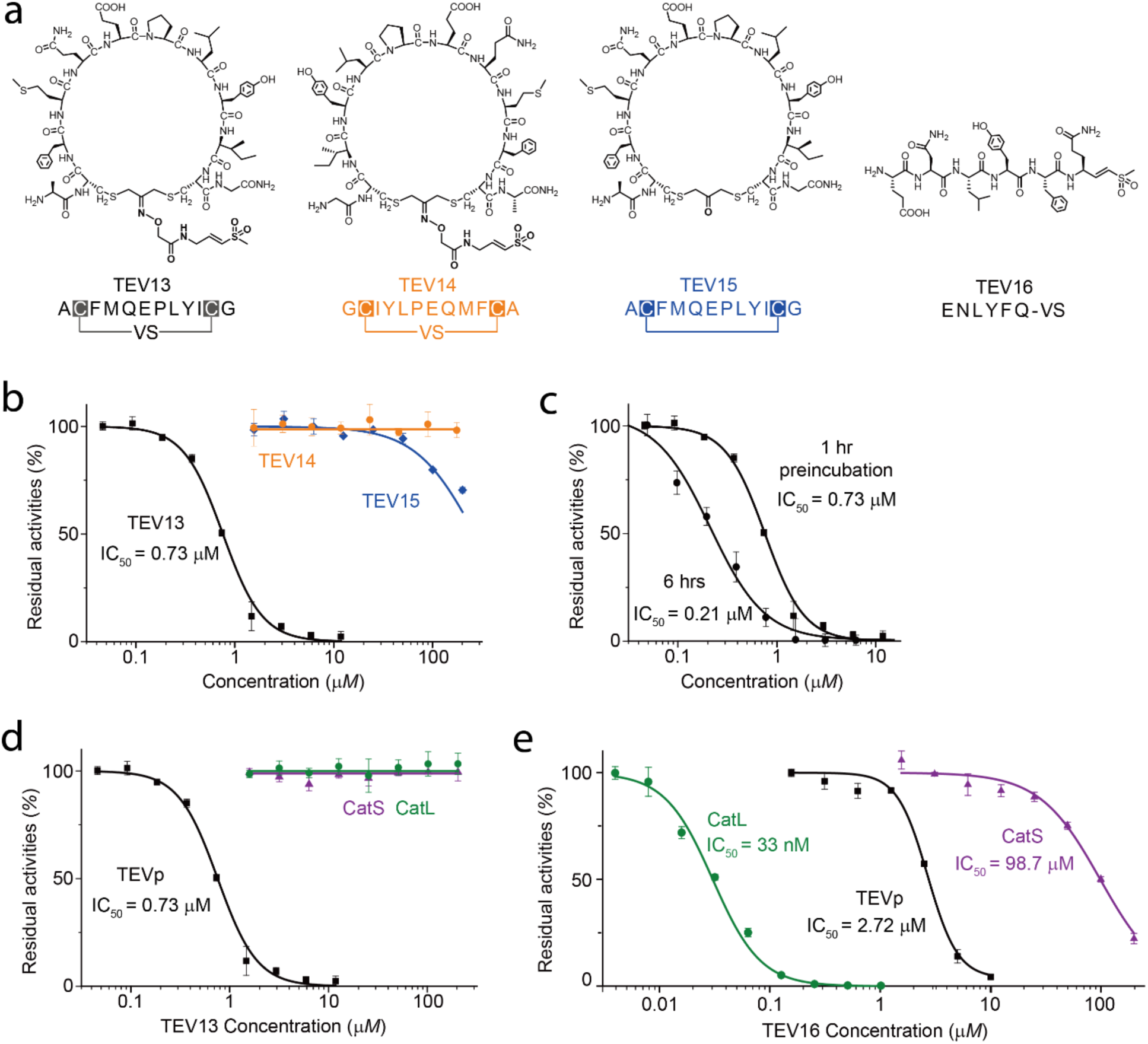
TEV13 is a potent and specific covalent inhibitor of TEV protease. a) Structures of the optimized cpVS inhibitor TEV13 compared to the scrambled isomer, TEV14, the cyclic peptide lacking the VS electrophile in the linker, TEV15, and the linear peptide VS synthesized based on the conserved substrate specificity of TEV. b) Dose response inhibition studies of the cyclized peptides using the recombinant TEV. Plots show residual enzyme activity over a range of inhibitor doses as measured by hydrolysis of a fluorogenic substrate. Enzyme was pre-treated with inhibitor at the indicated concentrations for 1 hour followed by addition of substrate and measurement of activity. IC_50_ values, when measurable are indicated. c) Time dependence of inhibition of TEV protease by TEV13. Recombinant enzyme was preincubated with a range of TEV13 concentrations for 1 hr or 6 hrs as indicated followed by measurement of residual activity using a fluorogenic substrate. Comparisons of the activity of (d) TEV13 and (e) TEV16 for TEV and the off-target cysteine proteases Cat S and Cat L. Inhibition studies were performed as in (b). n= 3 biologically independent wells (mean + s.d. is depicted).

Testing of the cyclic peptide controls for inhibition of recombinant TEV confirmed that the scrambled sequence (TEV14) had no measurable activity even at concentrations as high as 200 μM while the cyclic peptide lacking the VS warhead (TEV15) showed weak binding with less than 50% inhibition at concentrations over 200 μM (Fig. 4b). Furthermore, the potency of the optimized inhibitor TEV13 increased with increasing incubation time consistent with a covalent irreversible inhibition mechanism (Fig. 4c). We then tested the linear peptide VS, TEV16 and found that it was a relatively potent inhibitor with an IC_50_ value only a few times higher than the optimal cyclic peptide inhibitor TEV13 (Fig. 4d).

To determine the selectivity of TEV13 compared to the linear peptide TEV16, we tested both against other cysteine proteases. We chose sentrin-specific protease 1 (SENP1) because it has been successfully targeted with a peptide vinyl sulfone attached to a short linear peptide[38] or the full length folded SUMO protein domain[39]. Interestingly, neither the linear TEV16 nor the cyclic TEV13 was able to inhibit this protease at concentrations as high as 200 μM (Supplement Fig. 4). We then tested the inhibitors against two cysteine cathepsins (Cat S & Cat L) because both are relatively abundant and active proteases with broad substrate specificity profiles[40] that have been effectively targeted with peptide vinyl sulfones[41]. Importantly, we found that TEV13 failed to inhibit cathepsin S or L even when preincubated for 1 hour at 37 °C at concentrations over 200 μM (Fig. 4d). In stark contrast, the linear peptide VS TEV16 showed highly potent inhibition of cathepsin L with an IC_50_ value in the low nanomolar range and substantial inhibition of Cat S when used at high micromolar concentrations (Fig. 4e). Thus, even though the linear peptide that was rationally designed based on the highly conserved substrate specificity of TEV and is a reasonably good inhibitor of TEV, it is two orders of magnitude more potent for Cat L. This result highlights the fact that the phage screening approach enabled selection of binding elements that can selectively direct the vinyl sulfone to a target protease even when an off-target protease is efficiently inhibited by highly related linear peptide containing the same reactive electrophile.

### Generation of a selective ABP from phage selected inhibitors

One of the most effective ways to assess the selectivity of a covalent binding molecule is to make a fluorescent version for labeling complex mixtures containing a larger number of possible off-target proteins and other reactive nucleophiles[42]. We therefore conjugated a Cy5 fluorophore to the free N-terminus of TEV13, TEV14 and TEV16 to generate the corresponding fluorescent probes (Fig. 5a). We tested probe labeling of purified TEV protease in buffer as well as when added to total protein extracts from HEK293 and TG1 cells. We found that Cy5-TEV13 specifically labeled TEV protease both alone and when mixed with complex protein mixtures. Importantly, the corresponding scrambled peptide probe TEV14 showed no labeling of the target, consistent with its lack of activity for TEV (Fig. 5b). The linear peptide VS probe Cy5-TEV15 effectively labeled TEV both in buffer and in total protein extracts. However, the intensity of TEV labeling was reduced compared to Cy5-TEV13 and the probe also showed an increased amount of background labeling in the lysates, suggesting a decreased overall selectivity (Fig. 5c). Covalent binding of Cy5-TEV13 and Cy5-TEV14 were also concentration and incubation time dependent (Supplement Fig. 5). Furthermore, the TEV labeling by Cy5-TEV13 could be blocked by preincubation with the TEV13 inhibitor lacking the Cy5 label (Supplement Fig. 6). These results suggest that the TEV13 probe has an exceptionally high degree of specificity even in the presence of other cysteine proteases and reactive nucleophiles. To further assess selectivity of the probes, we performed labeling of RAW cell extracts with various concentrations of the probes at two different pH values. We chose RAW cells for their high levels of cysteine cathepsin expression[43]. The labeling results confirmed that the Cy5-TEV13 probe was highly selective with virtually no off-target background labeling even at the highest probe concentrations used at both pH 5.5 and 7.4 (Fig. 5d & Supplement Fig .7). In contrast, the linear peptide Cy5-TEV16 labeled multiple cathepsins even at the lowest probe concentration, and showed signs of labeling other off-target proteins indicated by increased background staining (Fig. 5D & Supplement Fig. 7). We found that the labeling efficacy of Cy5-TEV16 for cathepsin L was similar to the optimized cathepsin probe BMV109[18].

**Fig. 5.**
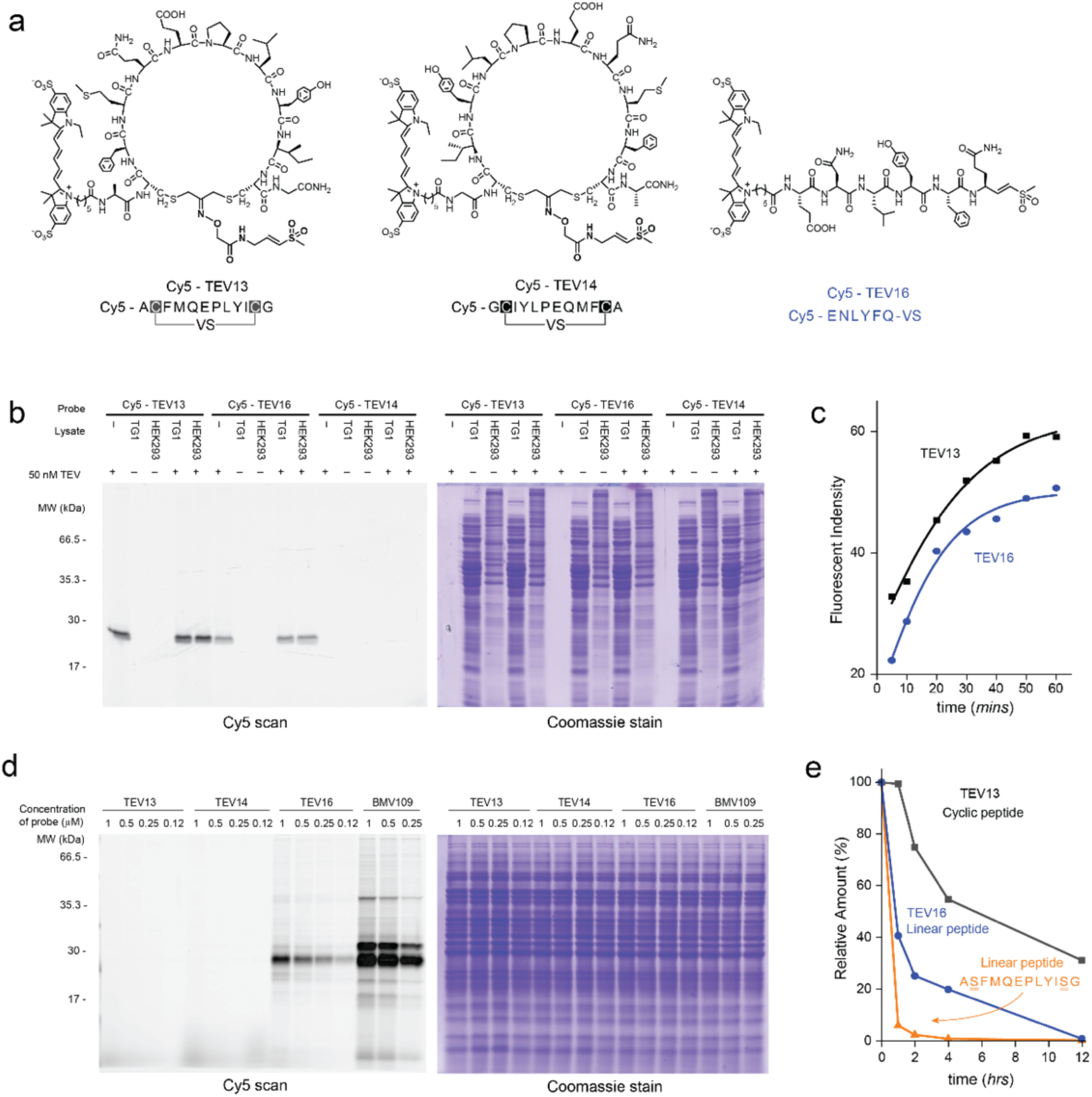
The Fluorescent cpABP Cy5-TEV13 specifically labels TEV protease in complex proteomic samples. a) Chemical structures of the Cy5 fluorophore labeled ABP versions of the cyclic peptides TEV13, TEV14 and the linear peptide-VS TEV16. b) Samples of enzyme buffer (-) or lysates from TG1 or HEK293 cells (4 μg total protein) were spiked with purified recombinant TEV protease (15 ng) or buffer as indicated, followed by addition of the indicated Cy5-labeled probes. Samples were incubated for 1 hr at 37 °C and then SDS PAGE sample buffer added followed by boiling. Samples were separated on SDS-PAGE gels followed by scanning for Cy5 fluorescence using a flatbed laser scanner (Left) and then stained with Coomassie brilliant blue (Right) to visualize total protein loading. c) Curves plotting the intensity of labeling of recombinant TEV protease (15 ng) in buffer over the range of indicated times. Samples were analyzed by SDS-PAGE analysis followed by scanning of the gel using flatbed laser scanner to quantify the intensity of labeled protease at each time point. d) Images of SDS-PAGE gels containing samples of the indicated total cellular lysates labeled for 1 hr with the indicated concentrations of each Cy5-labeled probe. The probe BMV-109 is a general ABP for cysteine cathepsins that was included for reference. The gels were processed as in (b). e) Plots of relative levels of each of the indicated cyclic and linear peptides after the indicated time of incubation in mouse plasma. Values were obtained by quantification of parent ions by mass spectrometry.

**Fig. 6.**
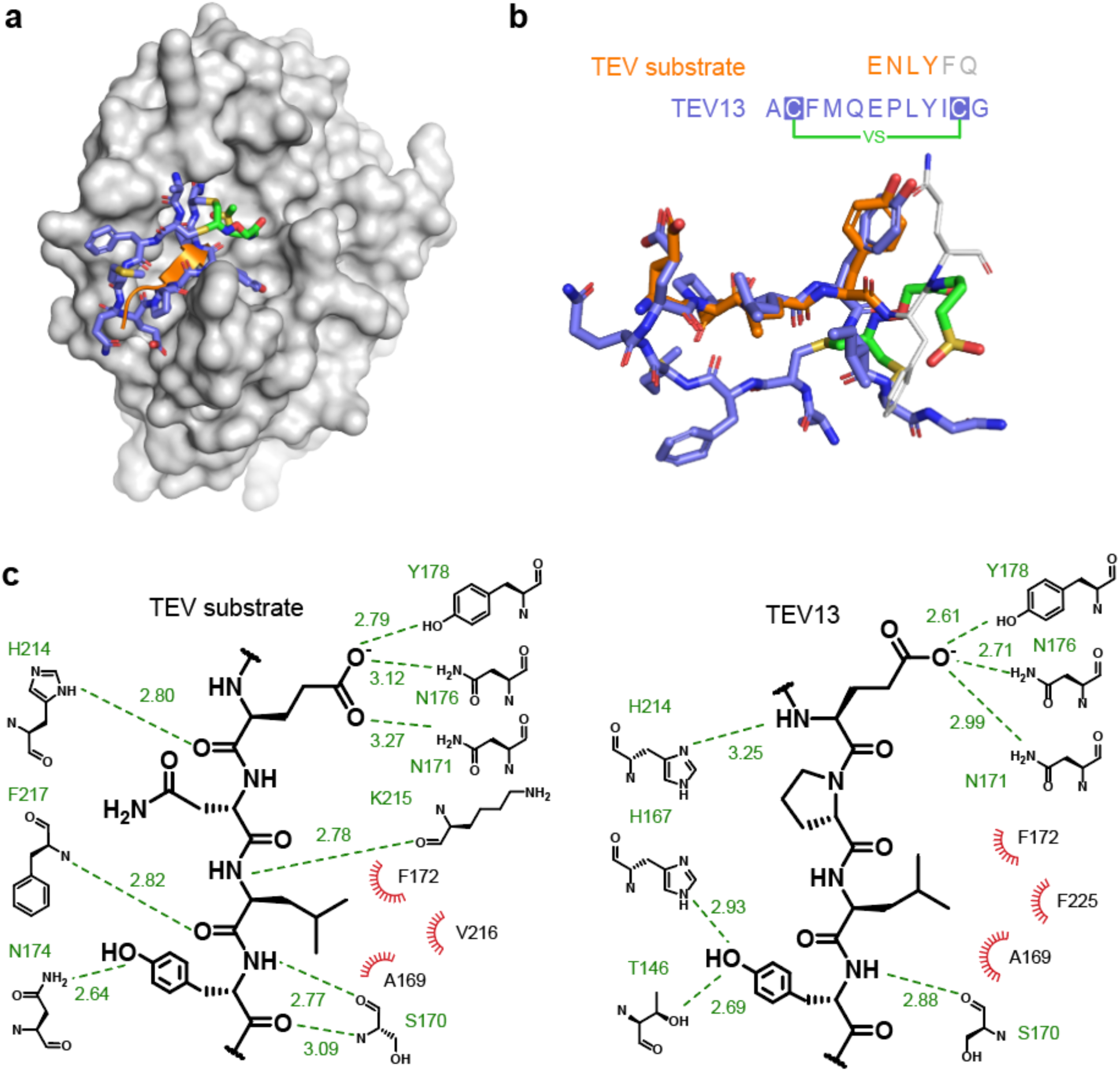
Molecular dynamic simulations to predict the interactions between TEV13 and TEV protease. a) The MD minimized complex structure of TEV13 (colored in blue) bound to TEV protease (colored in gray). The native substrate of TEV protease is shown as an orange ribbon as found in the reported crystal structure (PDB ID: 1LVM). b) Structure of the MD optimized TEV-bound TEV13 layered onto the reported X-ray crystal structure of the native peptide substrate ENLYFQ. The key residues P3-6 that overlap with residues in TEV13 are shown in orange. c) Extended diagrammatic representation of interactions between key residues on the native substrate and TEV13 and the active site region of the TEV protease. H-bond interacting partners are shown with green dash lines with distance indicated in Å. Hydrophobic interactions are depicted as red radiations.

### Plasma stability of cyclic peptide ABP

In addition to providing potentially improved target specificity by inducing protein-like folds, the cyclization of peptides also improves overall metabolic stability[44]. We therefore tested the serum stability of the optimal cyclic probe TEV13 and compared it to the linear peptide TEV16 as well as a linear version of the peptide portion of TEV13 in which the two cysteine residues are replaced with serine to prevent disulfide bond formation in the absence of the cyclization linker (Fig. 5e and Supplementary Fig. 8). After incubating TEV13 and the linear peptide analogs in mouse serum at 37 °C for 0, 1, 2, 4 and 12 hours, we preformed extraction of the peptides and HPLC analysis to quantify the amount of remaining parent molecules. These data confirmed that both linear peptides were degraded within the first hour, while the cyclic peptide remained fully stable for 1 hour and then only partially metabolized over the next 12 hours. Overall, these results confirm that use of a cyclic peptide backbone results in dramatically increased stability of inhibitors and probes.

### Interactions between cyclic peptide and TEV protease

To better understand how the cyclic peptide selected by phage screening is able to induce selective binding to the TEV target, we sought to analyze this protein-ligand interaction using structural information. We applied molecular dynamics (MD) to simulate the interactions between TEV13 and TEV protease using the reported high-resolution structure of TEV protease bound to a 6-residue substrate peptide as a template (ENLYF; PDB ID: 1LVM, chain C). While the power of MD calculations for *de novo* prediction of ligand-protein interactions is limited in some cases, there are number of reasons why MD simulations are informative for predicting the interactions between TEV protease and TEV13. Firstly, the covalent linkage between TEV13 and TEV protease largely defines the orientation in which the peptide ligand can bind to the target protein, thus greatly narrowing the chemical space to explorer and improving the probability of discovering relevant interactions. Secondly, the critical residues in the cyclic peptide portion of TEV13 show high similarity to the native TEV substrate sequence of EXLY<u>XQ-S</u>/G (where X can be any amino acid, underlined residues were not present in TEV13)[26]. This allows us to generate a more reliable initial conformation of TEV13 based on these key residues in the native substrate. To perform the MD simulations, we covalently linked the vinyl sulfone warhead of TEV13 to the active site cysteine of TEV protease to generate input files that were then loaded into the Assisted Model Building with Energy Refinement (AMBER) biomolecular simulation package[45]. The complex structure was subjected to MD simulation in explicit TIP3P solvent models with ff14SB and gaff parameter sets[46, 47]. After 40 ns of production simulation, the binding of TEV13 to TEV protease reached a constant and stable status (Supplementary movie). The energy of the entire system remained smooth in the simulation and the structure converged to a single low energy state for the complex (Supplementary Fig. 9). This predicted lowest energy structure shows TEV13 stably bound to the TEV protease in the primary substrate binding pocket (Fig. 6a). The side chains of residues Glu6, Leu8 and Tyr9 adopted a constant conformation similar to the native substrate peptide that involves strong interactions with TEV protease (Fig. 6b). In particular, Tyr9 in TEV13 adopted the same orientation as the P3 tyrosine in the TEV substrate peptide, forming a backbone H-bond with Ser170 and a hydroxyl group that interacts with residues of Asn174, T146 and H167 that form the S3 pocket of the TEV active site (Fig. 6c). Leu8, the same as P4 leucine in TEV substrate, located in the S4 hydrophobic pocket generated with Phe172, Val216, F225 and Ala169. Glu6 inherited the role of P6 glutamate by forming several H-bonds with Tyr178, Asn176, Asn171 and His214 located at the lower end of the substrate recognition epitope of TEV protease. Substitution of these three residues in the TEV substrate resulted in progressively decreased processing[48], which matched the result of our sequence-activity relationship (SAR) study (Fig. 3c). The MD results also indicated that Ile at position 10 contributed to the hydrophobic pocket formed by Val216 of TEV and Leu8 of TEV13. This could explain the preference of hydrophobic reside at the 10 position of the cyclic peptides. The flexible loop of the cyclic peptide in TEV13 composing residue 3-5 seems to only interact weekly with the N-terminal β-sheet 214-217 of TEV protease, which could explain the relatively narrow SAR profile that we observed for residue 3-5. However, the optimized MD structure supports the contribution of cyclization through loop 3-5 to increased entropy resulting in tight binding of TEV13 in the active site. Overall, MD simulations can make predictions about the optimized structure of the complex, and it confirms that stable and low energy confirmations can be found in which the cyclic peptide portion of the TEV13 molecule presents amino acid residues into established substrate binding pockets such that the reactive vinyl sulfone is positioned for covalent modification by the active site cysteine residue.

## Discussion

The recent advances in the applications of covalent activity-based probes combined with a revitalized interest in covalently binding drugs[1] suggest that there will be a need to find optimal ways to identify these types of chemically reactive ligands. ABPs have proven to be valuable tools for the study of many diverse classes of enzymes with applications ranging from identification of uncharacterized enzymes, to high-throughput screening to identification of new classes of small molecule inhibitors of therapeutically relevant targets[4]. They have also been used as imaging agents in both cell biology and in biomedical applications[49-51]. Furthermore, recent approval of therapeutic agents that irreversibly target enzymes such as protease and kinases, demonstrate the value of molecules capable of inducing a controlled covalent reaction with their target[52, 53]. The greatest challenge for the continued development of new and useful ABPs and covalent drugs is finding more effective and rapid ways to directly screen for selective covalent binding scaffolds. Currently, the most common way to develop new covalent ligands involves starting with a potent and selective inhibitor of a target and identifying optimal locations for placement of a reactive functional group and reporter tag. This often requires prior access to a potent lead molecule and a significant amount of medicinal chemistry efforts to identify an optimal covalent binder from that lead. Even when such efforts are successful, in many cases the resulting compounds still lack absolute specificity for the target. In order to obtain selectivity over closely related enzymes, it is necessary to test a large diversity of complex ligand scaffolds. However, this diversity and complexity is generally too great for standard chemical synthesis methods. Here we demonstrate a phage display method to build cyclized peptides containing a reactive electrophile directly on the surface of the phage particle. This allows both screening of libraries of potential covalent binding ligands that number in the billions and enables iterative screening after amplification of binders resulting in identification of sequences with high affinity and exceptional selectivity.

We chose to focus on cyclic peptides based on the fact that both cyclic and bicyclic structures can be generated directly from peptide sequences on an expressed phage coat protein[21, 31, 54]. Therefore, we could use the chemical linker that induces formation of the cyclic peptide as a way to introduce the electrophile for targeting of the active site nucleophile. Furthermore, cyclizing the peptide scaffold results in structures that are more rigid and protein-like than simple linear peptides[25]. We therefore reasoned that such cyclic peptide structures would have increased potential to direct selectivity of an ABP. Our results for the TEV protease confirm this hypothesis.

We chose TEV for our initial validation of the phage screening approach because it has a well-defined substrate specificity and thus is likely to contain defined binding pockets for substrate amino acids[26]. Furthermore, there are currently no reported potent and selective inhibitors of TEV that could easily be converted into ABPs for this protease. Our results show that we selected sequences in the cyclized portion of the peptides that contain residues that are part of the established substrate binding region of TEV. Interestingly, these sequences only contain some of the key anchor residues for TEV binding and all lack the key P1 glutamine residue that is essential for TEV recognition in a linear peptide sequence[35]. These data suggest that by screening our highly diverse set of billions of cyclic peptide sequences that we identified confirmations that can present key anchor residues to the protease in the absence of other key binding interactions. Molecular dynamic simulation helped to clarify the binding mode and will direct future efforts to find optimal linkers for use with other enzyme targets.

The TEV protease was also an ideal choice as a screening target because it is one of the relatively small number of examples of proteases with a single, highly well-defined extended `substrate recognition sequence. Therefore, it is an optimal candidate for development of selective ABPs by simple conversion of the known cleavage site into a linear peptide containing a reactive electrophile. Our results show that while the linear vinyl sulfone inhibitor containing optimal TEV substrate recognition sequence (TEV16) is in fact an effective inhibitor of TEV, it is significantly more potent inhibitor of cathepsin L. Consistent with increased activity towards an off-target cysteine protease, conversion to the Cy5 labeled version (Cy5-TEV16) showed that the resulting ABP, in addition to labeling cat L, also shows substantial cross-reactivity that results in overall high background signals when added to complex proteomic samples. These data highlight the fact that, even for highly selective proteases, short linear peptide sequences are often unable to induce sufficiently selective binding to a single target protease. By using the phage screening approach, it was possible to identify sequences from billions of possible ligands such that the identified hits are presented to the protease in a unique way that cannot be replicated for other, even closely related proteins.

The phage screening approach described here has the potential to be used broadly to identify covalently binding ligands for a diverse range of biomolecular targets. Our results suggest that the approach works well for proteases using a vinyl sulfone electrophile. It is likely that the same general linker could be used for screening against other classes of cysteine proteases but it could also possibly be used for other targets that contain a reactive cysteine residue. For example, there have been several recent reports of inhibitors that target cysteine residues in protein kinases[2] and proteomic profiling studies have begun to catalog all potential reactive cysteine residues in a cell[55]. This work has been complemented by other global studies of lysine reactivity which potentially identifies targets that would be amenable to screening for covalent ligands using this approach[56]. We are currently actively working to develop a serine reactive electrophile containing linker to enable phage screening for selective covalent ligands for diverse families of serine hydrolase enzymes that have been effectively isolated and labeled using the general fluorophosphonate ABPs[57].

One of the critical considerations for the success of the phage panning approach to identify selective covalent ligands is the choice of an optimal electrophilic warhead group. It is critical to use molecules that have relatively weak intrinsic reactivity towards free nucleophiles to avoid issues with non-specific covalent interactions with the phage. One of the primary benefits of using phage for selection is the ability to generate a large diversity of peptides on the phage surface. However, this diversity can also enhance the potential for peptides containing the reactive electrophile in the linker to bind targets in a non-specific way. Since screening involves selection and amplification, even a low level of non-specific binding can quickly dilute true binders of the active site nucleophile. We chose to use a vinyl sulfone for these initial studies because this electrophile is known to function as a weak electrophile when placed in the context of a peptide backbone[41]. Furthermore, vinyl sulfones are generally selective for thiol nucleophiles due to their propensity to undergo Michael addition reactions. Our results confirm that the vinyl sulfone is in fact an ideal electrophilic warhead for use in ABPs. We found that, of the phage clones that were enriched for covalent TEV binding, all contained sequences that had consensus binding residues known to be optimal for TEV. Our results using inhibitors and probes containing the same residues as the optimal TEV13 sequence but in scrambled order shows that even when anchor residues are present, if they are not oriented such that the can display the weak electrophile to the reactive nucleophile, there is no reaction.

In summary, we believe that our phage screening approach could be used to find covalent binding ligands for virtually any biomolecule target (protein, glycan, nucleotide, lipid, etc) that contains a slightly reactive nucleophile and the potential to interact with a macromolecular ligand. Thus, the covalent phage screening approach could greatly enhance the application of selective covalent ligands in cell biology, imaging and drug development.

## Acknowledgements

The authors thank the Vincent Coates Foundation Mass Spectrometry Laboratory at Stanford University for providing technical assistance with mass spectrometry. The authors also thank Dr. David Waugh for providing the TEV expression construct pDZ2087 and Dr. Christian Heinis for providing the phage library. The work was supported by The Swiss National Science Foundation Postdoc Mobility fellowship P2ELP3_155323 P300PB_164725 (to S.C.) and by funding from The National Institutes of Health grants R01 EB026285 and R01 EB026285 02S1 (to M.B,).

## Author contributions

M.B. and S.C. conceived the project and designed the experiments. S.C. performed the experiments. M.B. and S.C. analyzed the data and wrote the manuscript.

## Data availability

All data presented in this manuscript are available from the corresponding author upon reasonable request. The TEV protease expressing plasmid sequence is availability at GenBank with accession number MN480436. Synthetic methods and characterizations of cyclization linker and vinyl sulfone probes are available in the Supplementary Information.

## Materials and Methods

### General synthesis methods

Chromatographic separations were performed by manual flash chromatography unless otherwise specified. Silica gel 60 (Merck 70-230 mesh) was used for manual column chromatography. Merck F-254 thickness (0.25 mm) commercial plates were used for analytical TLC to follow the progress of reactions. Unless otherwise specified, ^1^H NMR spectra and ^13^C NMR spectra were obtained on a Varian Mercury 400 MHz console connected with an Oxford NMR AS400 Actively Shielded magnet, or a 500 MHz Varian Inova spectrometer at room temperature. Chemical shifts for ^1^H or ^13^C are given in ppm (δ) relative to tetramethylsilane (TMS) as an internal standard. Mass spectra (m/z) of chemical compounds were recorded on a Thermo Finnigan LTQ mass spectrometer, waters Acquity UPLC SQD2 system or Agilent 1260 Infinity LC connected with an InfinityLab MS detector. Reverse phase HPLC purifications were performed in 1260 infinity HPLC equipped with a semi-prep C-18 column (Phenomenex, 5 µm C18 100 Å, 250 × 10 mm, LC Column) eluted over a linear gradient from 95% solvent A (H_2_O, 0.1% TFA) to 100% solvent B (ACN, 0.1% TFA).

### Expression of biotinylated TEV protease with AviTag (Biotin-TEVp)

The biotinylated TEV protease (Biotin-TEVp) expression vector pMAL-HisTag-AviTag-3Cs-TEV was built by inserting the AviTag sequence into the pRK793 vector.^1^ TEV protease catalytic domain and vector backbone were amplified by polymerase chain reaction (PCR) with primers TEV_forwards (5’-GTTTTATTCCAGGGTCCTAGTGGTGGCGGTGGAGAAAGCTTGTTTAAGGGGC-3’) and TEV_reverse (5’-ACCTTGAAAATAAAGATTTTCTCCCC-3’). The AviTag sequence was amplified from pAviTag N-His Kan Vector (Ludigen, #49041-1) with primers AviTag-5 (5’-GAGAAAATCTTTATTTTCAAGGT CATCATCACCACCATCACGG-3’) and AviTag-3 (5’-AGGACCCTGGAATAAAACTTCTAAAGAGCCACCGGTAGGAGGAG-3’). The two PCR products were then ligated with Gibson Assembly (New England Biolabs, E2611S). The sequence of the final plasmid was verified with sanger sequencing (McLab, south San Francisco). The plasmid was transformed into Biotin XCell F’ cells for expression. An over-night pre-culture in 5 mL 2YT media containing 100 μg/mL Carbenicillin was then used for inoculating 500 mL 2YT median containing 100 μg/mL Carbenicillin and 50 μM biotin on the second day. After culturing at 37 °C until cell density reached OD_600_ of 0.4, 0.01 % arabinose and 1 mM IPTG was added to induce the expression of the BirA biotin transferase and the desired TEV protease. After further growing at 37 °C for another 4 hours, the cells of the expression culture were harvested by centrifugation and lysed in 50 mL 50 mM Tris 8.0, 100 mM KCl, 1 mM EDTA, 1 mM DTT, 0.01% Triton X100 buffer containing 10 mg lysozyme and 0.5 mg DNase with the help of sonication. After further incubation on ice for 30 mins, the insoluble cell debris was centrifuged at 10,000Xg for 20 mins. The clarified supernatant was pumped through a HisTrap™ HP 1mL column with a peristaltic pump (Rainin Dynamax RP-1) at a flow of 1 mL/min. After washing with 50 mL 50 mM Tris 8.0, 100 mM KCl, 1 mM EDTA, 1 mM DTT and 0.01 % Triton X100 buffer with 10 mM imidazole, the bound protein was eluted with an increasing imidazole gradient from 10 mM to 300 mM over 30 mins at a flow rate of 1 mL/min. The eluted protein fractions were combined and concentrated to 5 mL before being injected into a HiPrep 16/60 Sephacryl S-100 HR (GE healthcare) for purification. The purified fractions of TEV protease were analyzed by SDS-PAGE and fractions containing the enzyme were combined and stored in 50 mM Tris 8.0, 0.5 mM EDTA and 1 mM DTT at −80 °C.

### Peptide-D1-D2 fusion protein expression

A previously reported plasmid^2^ (pET28b-C7C-D1D2) was used for expressing a peptide (ACGSGSGSGCG) containing two cysteine residues fused to the phage D1D2 protein (ACGSGSGSGCG-D1D2). The expression and purification procedures are similar to the expression procedure described above except the following changes were made: BL21(DE3) cells were used for expression at 30 °C for 14 hours. The final protein was eluted with the buffer R (20 mM NH_4_HCO_3_, 5 mM EDTA, pH 8.0 with 1 mM TCEP).

### Modification of the peptide-D1-D2 fusion protein

Excess TCEP in the D1D2 fusion protein was removed by exchanging the protein buffer to buffer R without TCEP using a PD-10 column (GE Healthcare). The concentration of the protein was determined using a Nanodrop spectrometer based on absorption at 280 nm. The DCA linker (300 μM) was added to 2 μM fusion protein in buffer R and incubated at 30 °C 2 hours or 42 °C for 1 hour before being analyzed using a Waters ACQUITY UPLC equipped with a C8 column and SQ Detector 2. The acquired mass of the protein was deconvoluted using MassLynx software.

**Supplementary Scheme 1.**
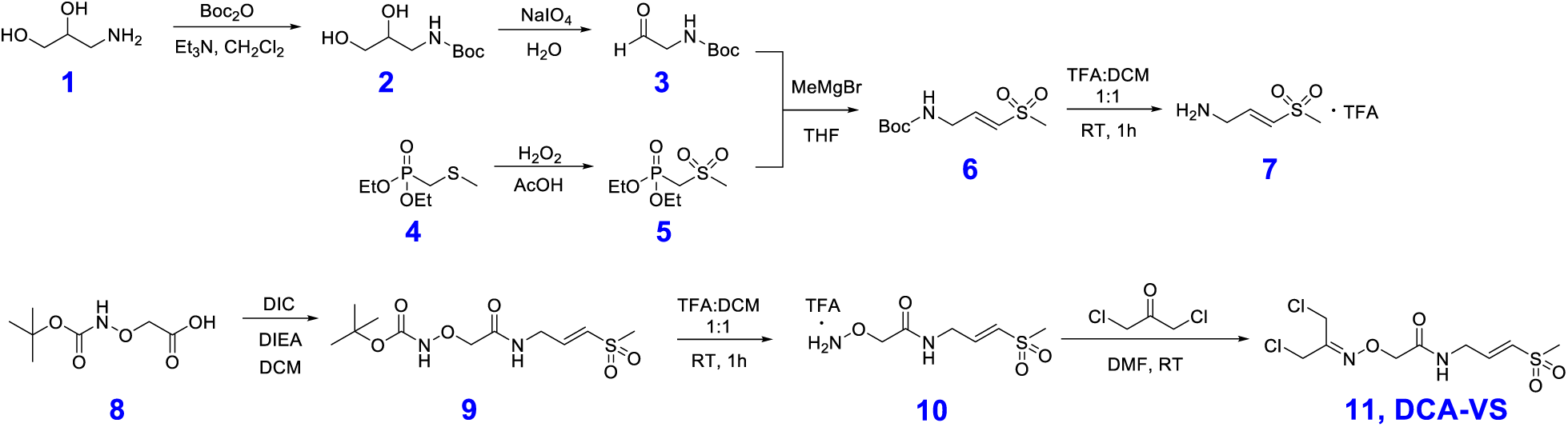
Synthetic strategy of DCA-VS, derivatizing 1,3-Dichloroacetone with a vinyl sulfone cysteine reactive warhead.

### DCA-VS linker synthesis

#### tert-butyl (2,3-dihydroxypropyl)carbamate (**2**)

Triethylamine (4.11 mL, 29.5 mmol) was added dropwisely to a solution of 1-aminopropane-2,3-diol (23.2 g, 255 mmol) in methanol at room temperature, following by slow addition of a solution of di-tert-butyl dicarbonate (66.8 g, 306 mmol) in dichloromethane (200mL). The reaction mixture was further stirred at room temperature for 3 hours. After removing the solvent, the oil-like residue was purified using a silica column and eluted with dichloromethane/methanol to give 40 g of the desired product as an oil (82% yield). ^1^H NMR (500 MHz, Chloroform-*d*) δ 5.02 (sb, 1H), 3.89 – 3.72 (m, 1H), 3.68 – 3.54 (m, 2H), 3.46 – 3.17 (m, 2H), 2.81 (sb, 2H), 1.47 (s, 9H).

#### tert-butyl (2-oxoethyl)carbamate (**3**)

Sodium periodate (53.78 g, 251 mmol) was added in portions to a tert-butyl 2,3-dihydroxypropylcarbamate (40 g, 209 mmol) solution in water (300 mL). After stirring at room temperature for 2 hours, the resulting solids were removed by filtration, and the aqueous fraction was extracted 6 times with dichloromethane. The combined organic fractions were washed with brine and dried over magnesium sulfate. After filtration, the dichloromethane fraction was dried in a rotary evaporator and used directly for the next synthesis step (crude yield 85%). ^1^H NMR (500 MHz, Chloroform-*d*) δ 9.68 (s, 1H), 5.21 (sb, 1H), 4.10 (d, *J* = 5.1 Hz, 2H), 1.48 (s, 9H).

#### diethyl ((methylsulfonyl)methyl)phosphonate (**5**)

To a solution of diethyl (methylthio)methylphosphonate (10 g, 50.5 mmol) in acetic acid (34.6 mL, 605 mmol) was slowly added 50% hydrogen peroxide in water (7.4 mL, 260 mmol). Following completion of addition, the reaction mixture was heated to 70 °C for 0.5 hour. After removing the solvent in a rotary evaporator, the oily residue was diluted with water and extracted six times with dichloromethane. The organic fraction was further washed with saturated sodium bicarbonate in water, brine and dried over anhydrous magnesium sulfate. After filtration, the dichloromethane fraction was further dried in a rotary evaporator. The resulting white solid was recrystallized in a mixture of ethyl acetate / hexane to give 9.3 g of a colorless crystalline solid (yield 80%). ^1^H NMR (500 MHz, Chloroform-*d*) δ 4.57 – 4.11 (m, 4H), 3.62 (d, *J* = 16.4 Hz, 2H), 3.24 (s, 3H), 1.41 (t, *J* = 7.1 Hz, 6H).

#### tert-butyl (E)-(3-(methylsulfonyl)allyl)carbamate (**6**)

In dry flask under argon, methylmagnesium bromide solution (3M; 13 mL, 39 mmol) in diethyl ether was added slowly through syringe to the solution of diethyl ((methylsulfonyl)methyl)phosphonate (8.95 g, 38.9 mmol) in dry tetrahydrofuran (100 mL). After stirring at room temperature for 15 mins, tert-butyl (2-oxoethyl)carbamate (6.8 g, 42.8 mmol) solution in dry tetrahydrofuran (100 mL) was added very slowly. After addition, the solution was further refluxed for 2.5 hours. To stop the reaction, 20 mL ammonium chloride saturation solution in water was added. Three times 40 mL diethyl ether was added to extract the product. After drying with magnesium sulfate, the diethyl ether fraction was dried in a rotary evaporator and the crude product purified using a silica column. The desired product was eluted with Hexane and ethyl acetate at a ratio of 3:1 (6.6 g; yield 72 %). ^1^H NMR (500 MHz, Chloroform-*d*) δ 6.94 (dt, *J* = 15.7, 4.8 Hz, 1H), 5.97 (dt, *J* = 15.8, 1.9 Hz, 1H), 4.73 (s, 1H), 3.95 (t, *J* = 5.6 Hz, 2H), 3.76 (s, 3H), 1.48 (s, 9H). ^13^C NMR (126 MHz, Chloroform-*d*) δ 166.55, 145.26, 120.71, 79.73, 60.37, 51.58, 41.28, 28.30.

#### tert-butyl (E)-(2-((3-(methylsulfonyl)allyl)amino)-2-oxoethoxy)carbamate (**9**)

A solution of tert-butyl (E)-(3-(methylsulfonyl)allyl)carbamate (80 mg, 0.34 mmol) in 50 % trifluoroacetic acid with dichloromethane was stirred at room temperature for 1 hour before the solvent was removed under vacuum and toluene added twice to co-evaporate the TFA. The remaining oily residue was triturated with diethyl ether to produce a brownish solid (50 mg, 0.216 mmol). Then 2-(((tert-butoxycarbonyl)amino)oxy)acetic acid (50 mg, 0.262 mmol) was added followed by *N,N’*-Diisopropylcarbodiimide (27.2 mg, 0.216 mmol) and *N,N*-Diisopropylethylamine (38 uL, 0.218 mmol). After stirring at room temperature overnight, the insoluble solids were filtered off and the dichloromethane fraction was purified directly by silica chromatography to produce 42 mg of product, for a 63% yield. ^1^H NMR (500 MHz, Chloroform-*d*) δ 8.68 (sb, 1H), 7.81 (s, 1H), 6.92 (dt, *J* = 15.2, 4.3 Hz, 1H), 6.55 (dt, *J* = 15.2, 2.0 Hz, 1H), 4.37 (s, 2H), 4.18 (ddd, *J* = 6.2, 4.3, 2.0 Hz, 2H), 2.93 (s, 3H), 1.47 (s, 9H). ^13^C NMR (126 MHz, Chloroform-*d*) δ 169.32, 158.18, 143.91, 129.88, 105.00, 83.69, 42.84, 38.88, 28.06.

(E)-2-(((1,3-dichloropropan-2-ylidene)amino)oxy)-N-(3-(methylsulfonyl)allyl)acetamide (DCA-VS) (**11**) Tert-butyl (E)-(2-((3-(methylsulfonyl)allyl)amino)-2-oxoethoxy)carbamate (42 mg, 0.136 mmol) was placed in 1 mL of a 50 % trifluoroacetic acid solution in dichloromethane at room temperature for 1 hour. The solvent was removed under vacuum. Toluene was added and dried twice to remove the residual trifluoroacetic acid. The resulting alkoxyl amine was mixed with 1,3-Dichloroacetone (25 mg, 0.198 mmol) in 1 mL of dimethylformamide. The mixture was stored at room temperature overnight. After removing dimethylformamide, the residue was purified by silica chromatography and the desired product was eluted with 3% methanol in dichloromethane to produce 26 mg product for a 60 % yield. ^1^H NMR (400 MHz, Chloroform-*d*) δ 8.01 (s, 1H), 6.90 (dt, *J* = 15.2, 4.4 Hz, 1H), 6.46 (d, *J* = 15.2 Hz, 1H), 4.71 (s, 2H), 4.35 (s, 2H), 4.27 (s, 2H), 4.19 (ddd, *J* = 6.4, 4.5, 1.9 Hz, 2H), 2.94 (s, 3H). ^13^C NMR (101 MHz, cdcl_3_) δ 168.98, 154.30, 143.64, 130.22, 73.65, 42.84, 41.78, 38.84, 32.79. MS (ESI): calcd for C_9_H_15_Cl_2_N_2_O_4_S^+^: 317.0; found: 317.0.

**Supplementary Scheme 2.**
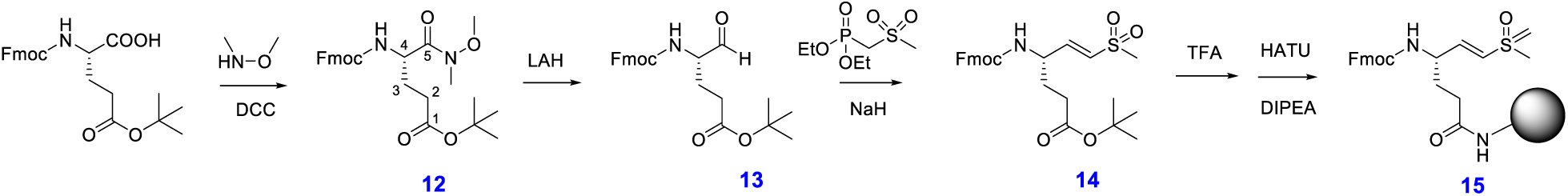
Synthetic strategy of Fmoc-glu-vinyl sulfone for preparing peptide-vinyl sulfone probes with a P1 glutamine by solid phase peptide synthesis.

#### Fmoc-Glu(OtBu)-dimethyl hydroxyl (Weinreb) amide (***12***)

Fmoc-Glu(OtBu)-OH (10.34 g, 24.3 mmol), HOBT (3.6g, 26.7 mmol) and dicyclohexylcarbodiimide (DCC; 5.5g, 26.7 mmol) were dissolved in DMF. After stirring 30 minutes at room temperature a solid formed and was removed by filtration. *N,O*-dimethyl hydroxyl amine (2.84g, 29.2 mmol) was added as a solid to the filtered reaction mixture along with triethyl amine (2.94g, 29.2 mmol). The reaction was stirred for an additional 12 hours and then concentrated by rotary evaporation of the solvent. The crude oil was then dissolved in ethyl acetate and extracted with three portions each of saturated sodium bicarbonate, 0.1N HCl and brine. The organic layer was dried over magnesium sulfate and concentrated to dryness. This crude preparation was used in subsequent reactions without further purification. ^1^H NMR (400 MHz, Chloroform-*d*) δ 7.76 (d, *J* = 7.4 Hz, 2H), 7.64 – 7.56 (m, 2H), 7.40 (t, *J* = 7.4 Hz, 2H), 7.36 – 7.27 (m, 2H), 5.60 (d, *J* = 8.8 Hz, 1H), 4.36 (d, *J* = 7.3 Hz, 2H), 4.21 (t, *J* = 7.2 Hz, 1H), 3.78 (s, 3H), 3.21 (s, 3H), 2.38 – 2.29 (m, 2H), 2.16 – 1.81 (m, 2H), 1.44 (s, 9H).

#### Fmoc-Glu(OtBu)-H (***13***)

Crude 12 (11 g, 23.5 mmol) was dissolved in anhydrous ethyl ether and stirred on ice under a positive argon flow. Lithium aluminum hydride (0.891 g, 23.5 mmol) was slowly added leading to gas evolution. The reaction was stirred for an additional 20 minutes and then quenched by the addition of potassium hydrogen sulfate (6.67g, 49 mmol). The quenched reaction was stirred on ice for 20 minutes and then at room temperature for an additional 30 minutes. The mixture was extracted three times with ethyl acetate and the combined organics washed with three portions of 0.1M HCl, saturated sodium carbonate and brine. The organic phase was concentrated to an oil. The product was purified by flash chromatography in hexanes:ethyl acetate (2:1, v:v). Yield (1 g, 10.4%). ^1^H NMR (400 MHz, Chloroform-*d*) δ 9.59 (s, 1H), 7.77 (d, *J* = 7.8 Hz, 2H), 7.60 (d, *J* = 7.5 Hz, 2H), 7.41 (t, *J* = 7.5 Hz, 2H), 7.32 (t, *J* = 7.5 Hz, 2H), 5.61 – 5.51 (m, 1H), 4.50 – 4.38 (m, 2H), 4.22 (t, 1H), 2.44 – 2.28 (m, 2H), 1.98 – 1.85 (m, 2H), 1.44 (s, 9H).

#### Fmoc-Glu(OtBu)-vinyl sulfone (***14***)

Diethyl (methylthiomethyl) phosphonate was oxidized to the corresponding phosphonate sulfone using hydrogen peroxide in acetic acid as described above (**5**). Diethyl phosphonate sulfone (96 mg, 0.417 mmol) was dissolved in anhydrous THF under argon. Methylmagnesium bromide solution (3 M; 139 μL, 0.417 mmol) in diethyl ether was added and the reaction and subsequently stirred for 30 minutes at room temperature. Pure **13** (170 mg, 0.416 mmol) was dissolved in anhydrous THF under argon and added to the stirring reaction by syringe. The reaction was then allowed to stir for 30 minutes, quenched with water and the resulting aqueous phase washed three times with dichloromethane. The organic phases were combined, dried over MgSO_4_ and evaporated to dryness. The crude oil was purified by flash chromatography over silica gel in hexane/ethyl acetate (1.5:1 v:v). Yield (168 mg, 8.7 mmol, 34.6%). ^1^H NMR (500 MHz, Chloroform-*d*) δ 7.81 (d, *J* = 7.5 Hz, 2H), 7.62 (s, 2H), 7.48 – 7.41 (m, 2H), 7.40 – 7.34 (m, 2H), 6.84 (dd, *J* = 14.6, 4.9 Hz, 1H), 6.46 (d, *J* = 15.0 Hz, 1H), 5.26 (d, *J* = 8.0 Hz, 1H), 4.49 (d, 2H), 4.23 (t, *J* = 6.7 Hz, 1H), 2.96 (s, 3H), 2.46 – 2.28 (m, 2H), 2.04 – 1.83 (m, 2H), 1.48 (s, 9H).

#### Fmoc-Glu(Rink amide resin)-vinyl sulfone (***15***)

Pure 14 (17 mg, 35 μmol) was dissolved in methylene chloride (0.5 ml) and an equal volume of anhydrous trifluoracetic acid was added. The reaction was stirred for 1 hours and quenched by the addition of toluene (1 mL). The reaction was concentrated by rotary evaporation, then re-suspended in toluene and dried twice. The crude product was then coupled to Rink amide resin (60 mg, 0.35 mmol/g load, 21 μmol) in DMF (1 mL) using HATU (13.3 mg, 35 μmol) and DIEA (11 μL, 105 μmol) for 2 hours at room temperature. The resin was washed with DMF and peptide chain extension was achieved with standard solid phase peptide synthesis.

### Phage panning

Phage harboring the cysteine rich peptide library (CX_8_C, X is any 20 canonical amino acid, C is cysteine) is a generous gift from Dr. Christian Heinis (EPFL, Lausanne). The phage peptide library in TG1 *E*.*coli* bacteria (Lucigen, # 60502-1) was thawed from stock and used for inoculating 1 L of 2YT media containing 30 μg/mL chloramphenicol. Phage were produced at 30 °C with shaking at 250 rpm for 16 hours. On the second day, the media rich with secreted phage was separated from the host bacteria by spinning at 8,500 rpm for 30 mins and the supernatant was mixed with 250 mL 20 % PEG 6000, 2.5 M sodium chloride solution. After being incubated on ice for another hour, the precipitated phage was spun at 9,000 rpm for 45 min. The resulting phage pellet was then dissolved in 10 mL buffer R (20 mM ammonium bicarbonate pH 8.0, 5 mM EDTA) and reduced with 1 mM TCEP at 42 °C for 1 hour. After removing the excess TCEP by filtration, 16 mL buffer R was added and 4 mL linker (DCA-VS; 1.5 mM) in acetonitrile was added to modify the phage at 30 °C for 2 hours. Then 5 mL 20 % PEG 6000, 2.5 M sodium chloride solution was added to precipitate the phage and resuspended in Buffer TEV (50 mM Tris 8.0, 100 mM potassium chloride, 1 mM EDTA and 1 mM DDT) supplemented with 1 % BSA and 0.1 % TX100 for blocking at room temperature for 0.5 hour. Biotin-TEV protease (10 μg) was incubated with 50 μL hydrophilic streptavidin magnetic beads (New England Biolabs, S1421S) for half hour and washed 5 times with TEV buffer supplemented with 1 % BSA and 0.1% TX100. After being blocked in the same buffer for half hour, the phage library was mixed with the magnetic beads and incubated at room temperature for different amounts of time, depending on the round of screening being performed. After washing 10 times with PBS buffer, the magnetic beads were incubated with 100 μL guanidine chloride in PBS for 5 mins and then eluted with 300 μL 3C protease at 30 °C for 1 hour to release the covalently bound phage from the solid support. The released phage were used to infect 45 mL TG1 bacteria at OD_600_ of 0.4 for 1.5 hours in a 37 °C incubator without shaking. Afterwards the TG1 cells were pelleted at 3,000 rpm for 15 min and plated on two 15 cm 2YT agar plate with 30 μg/mL chloramphenicol antibiotic. The agar plate was then incubated at 37 °C overnight to allow the infected TG1 cells to expand. On the second day, TG1 cells were scrapped off the agar plates with 5 mL 2YT median, mixed with 5 mL 50 % glycerol and stored at −80 °C. This glycerol stock was then used for preparing plasmid for sequencing or generating phage for next round of phage panning.

### Phage infectivity measurement

To the volume of 1 mL PBS containing 0, 1, 2, 3, 4, 5 and 6 M guanidine chloride, 5 μL of phage stock solution was added and incubated at room temperature for 5 mins. To the volume of 1 mL of TEV buffer containing 0-25 ng/mL TEV protease, 5 μL of phage stock solution was added and incubated at room temperature for 0.5 hour. The solutions at each condition (20 μL) was taken to perform 12 serial dilutions (10-fold) with 180 μL 2YT medium. Each dilution (20 μL) was added to 180 μL exponentially growing TG1 cells (OD_600_ = 0.4) and incubated for 90 minutes at 37 °C. 10 μL of the infected TG1 cells were then spotted onto 2YT agar plates containing 30 μg/ml chloramphenicol. The number of colonies was counted the next day and the number of infective phage was calculated.

### Peptide modification

All peptides were prepared using standard solid-phase peptide synthesis on a Syro II automated parallel peptide synthesizer (Biotage, Charlotte, NC USA) using standard 9-fluorenylmethoxycarbonyl (FMOC) chemistry protocols and Rink Amide AM resin support (0.02 mmol scale). Fmoc groups were removed using 300 μL of a 20% (v/v) solution of piperidine in DMF and amino acid coupling was carried out following a double coupling strategy with a 4-fold excess of Fmoc-amino acids (0.5 M solution in DMF). Coupling was achieved using a 1/1/1.5 ratio of amino acid/HBTU/DIEA in DMF. The deprotection and coupling times were 5 and 30 minutes, respectively. Six times 600 μL DMF washes were performed between deprotection and coupling steps. The peptide-loaded resin was treated with cleavage cocktail K (90/2.5/2.5/2.5/2.5 (v/v) mixture of TFA/thioanisole/water/phenol/EDT) for 2 h, which simultaneously cleaved the peptide from the resin and removed all side-chain protecting groups. The cleaved peptides were subsequently isolated by precipitation in cold diethyl ether followed by centrifugation. The precipitated peptides were resuspended in diethyl ether and centrifuged (3 times). Finally, the peptides were dissolved in degassed MilliQ water and lyophilized.

To perform peptide cyclization, 10 mg of the crude peptide was dissolved in 9 mL degassed solvent mixture of 20 % acetonitrile and 80 % water resulting in a peptide concentration of about 1 mM, followed by addition of 3.5 mg DCA-VS linker in 0.5 mL acetonitrile and 0.5 mL 1 M NH_4_HCO_3_ in water. After incubation at 30 °C for 1 hour, the reaction mixture was quenched with 50 μL TFA and lyophilized to driness. For purification, the modified peptides were dissolved in 400 μL water and injected onto a 1260 infinity HPLC (Agilent technologies, USA) equipped with a semi-prep C-18 column (Phenomenex, 5 µm C18 100 Å, 250 × 10 mm, LC Column), and separated over a linear gradient from 95 % solvent A (H_2_O, 0.1% TFA) to 100% solvent B (ACN, 0.1% TFA) in 15 mins. The mass of peptides was measured using a Bruker Maldi Microflex LRF, and HPLC fractions were lyophilized and resulting solids dissolved in DMSO to a stock concentration of 10 mM and stored at −20 °C.

### TEV protease expression

The TEV protease conjugated with a N-terminus six Histidine tag was expressed in *E. coli* strain BL21(DE3) (Lucigen, #60401-1) using the cytoplasmic expression plasmid pDZ2087 (precious gift from Dr. David Waugh). BL21(DE3) harboring pDZ2087 was used to inoculate 5 mL 2YT media with 100 μg/mL ampicillin and diluted with 500 mL 2YT media containing 100 μg/mL ampicillin for expression on the second day. The culture was shaked at 250 rpm at 37 °C until the density of the bacteria reached exponential phase (OD_600_ = 0.5) when 1 mM IPTG was added to induce the expression of TEV. After shaking at 30 °C 250 rpm for 8 hours, the bacteria were harvested by spinning at 8000 rpm for 15min. The purification procedure is the same as the one used for purifying biotin-TEVp described above.

### Fluorogenic substrate synthesis

The TEV substrate peptide (ENLYFQGK, 20 μmol) was synthesize by standard Fmoc based solid phase peptide synthesis on rink amide resin (CHEM-IMPEX INT’L INC., #02900, 0.33 mmol/g load) as described above. After deprotecting Fmoc from the N-terminus of the synthesized peptide, cy5 acid (14 mg, 22 μmol), HATU (12 mg, 31.5 μmol) and DIPEA (18 μL, 103 μmol) were added to the washed resin for coupling cy5 to the N-terminus of the synthesized TEV peptide for 2 hours at room temperature. After releasing and deprotecting the peptide with cleavage cocktail K, the cy5 labeled peptide was purified in HPLC and lyophilized to dryness. MS (ESI) negative: calculated for M-H^+^: 1620.7; found: 1620.4. Cy5-ENLYFQGK-NH_2_ (10 mg, 6.2 μmol), QSY21 (7 mg, 7.5 μmol) and DIPEA (6.5 μL, 37.4 μmol) were mixed in 1 mL DMSO and incubated at room temperature overnight. On the next day, the reaction mixture was injected into a RP-HPLC connected with a semi-prep C18 column to purify the Cy5-ENLYFQGK(QSY21)-NH_2_ as described above. MS (ESI) negative: calculated for M-2H^+^: 1221.4; found: 1221.4.

### TEV inhibition assay

The inhibitory activity of bicyclic peptides was determined by incubating TEV protease (50 nM) with different concentrations of inhibitor and quantification of the residual activity using the fluorogenic substrate synthesized as described above (50 μM, Cy5-ENLYFQGK(QSY21)-NH_2_). Peptide inhibitors and TEV protease were pre-incubated at 30 °C for 1 hour, and then the enzymatic assays were performed at 30 °C in buffer containing 50 mM Tris-Cl, pH 8.0, 100 mM KCl, 1 mM EDTA 1 mM DTT and 0.01% TX100. The TEV activity was measured by monitoring the change in fluorescence intensity during one hour using a Cytation 3 Multi-Mode Reader (excitation at 640 nm, emission recorded at 670 nm; BioTek Instruments, Inc, Winooski, VT, USA). Calculations were done with OriginPro 2019 software (OriginLab Corporation). Sigmoidal curves were fitted to the data using the following dose response equation wherein x = peptide concentration, y = % activity of reaction without peptide, A1 = 100%, A2 = 0%, p = 1. IC_50_ values were derived from the fitted curve.

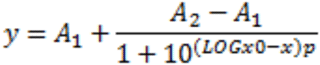

### Synthesis of SENP1 substrate Ac-QTGG-AFC

POCl_3_ (400 µL, 4.29 mmol) was slowly added to the solution of 7-amino-4-trifluoromethylcoumarine (AFC, 500 mg, 2.18 mmol) and Boc-Gly-OH (381 mg, 2.18 mmol) in pyridine (10 mL) at −15 °C under argon and the resulting mixture was further stirred at −15 °C for half hour. The reaction was then quenched with water and extracted with ethyl acetate. The organic phase was combined and washed with 0.1 M HCl, water, and brine. After dried over magnesium sulfate, the organic phase was concentrated in a rotary evaporator. The resulting crude material was purified by silica column to purify the final Boc-G-AFC as white solid (630 mg, 74% yield). After Boc deprotection with a mixture of TFA:DCM (1:1, 10mL) at room temperature for 1 hour, the reaction mixture was dried under a rotary evaporator and injected into a CombiFlash Companion Flash Chromatography System equipped with a RediSep ^®^ Rf Reversed-phase C18 column (Teledyne, Thousand Oaks, CA, USA) for purification. H_2_N-G-AFC was isolated as a white solid (550 mg, 88%). MS (ESI): calcd for M+H^+^: 287.2; found: 287.1.

Ac-Q(trt)T(t-Bu)G-OH was synthesized on chlorotrityl resin as described above and cleaved with 20% hexafluoroisopropanol in DCM 3×5 mins to keep the peptide fully protected. MS (ESI): calcd for M+H^+^: 645.3; found: 644.9. Ac-Q(trt)T(t-Bu)G-OH, H_2_N-G-AFC, DIC and DIPEA was mixed in DCM and stirred at room temperature overnight. The next morning, the generated white solid was removed by filtration and the reaction mixture was dried under vacuum and deprotected with 1 mL TFA:TIPS:water (95:2.3:2.5) for 2 hours at room temperature. After removing the solvent, the residue was dissolved in DMSO and injected onto an HPLC for purification. Fractions with the correct MS were combined and dried in a lyophilizer to give white solid. MS (ESI): calcd for M+H^+^: 615.2; found: 615.3.

### Cy5 labeled TEV13 peptide

Cy5-NHS (1 mg, 1.35 μmol) and DIPEA (1 μL, 5.75 μmol) was added to a solution of the TEV13 peptide cyclized with DCA-VS (10 mM, 50 μL) in DMSO. The reaction mixture was incubated at room temperature overnight and injected into HPLC for purification as described above. The fractions with the desired product were combined and lyophilized to dryness. MS (MALDI): calcd for M+H^+^: 2276.8; found: 2276.5.

### Labeling with Cy5-TEV13-DCA-VS

HEK293 cells from one 10 cm petri dish or TG1 *E*.*coli* grown in 5 mL 2YT media were individually resuspended in 1 mL TEV buffer (50 mM Tris 8, 100 mM KCl, 1 mM EDTA and 1 mM DTT) with 0.2% Triton X100. The cells were transferred to 2.0 mL O-ring tubes and mixed with 100 μL 0.1mm glass beads (Biospec Products, Bartlesville, OK, USA). Tubes were then placed in a prechilled aluminum vial rack and cell samples were lysed in a Mini-BeadBeater-96 (Biospec Prod) three times (30 s with 2 min resting on ice in between). After centrifugation at 13,000Xg, at 4 °C for 5 min, 50 μL of the cleared supernatant was mixed with 50 μL TEV buffer with/without 100 nM TEV protease for labeling with 1 μM Cy5-TEV13-DCA-VS at 30 °C for 1 hour. Then SDS PAGE loading buffer (40 μL) was added and the mixtures were heated to 95 °C for 5 mins. Samples from each condition (15 μL) were separated by 12% SDS-PAGE. Cy5 fluorescence was detected by scanning on a Typhoon 9410 variable mode imager (GE Healthcare, λ_ex_=633 nm, 670 BP30 filter). The fluorescent intensity of the labeled proteins at expected size was analyzed with LI-COR Image Studio Lite software, and plotted with OriginPro 2019 software (OriginLab Corporation). Afterwards, the gel was stained with Coomassie blue to visualize total protein loads.

### Ellman’s test of free cysteine in TEV13

Free cysteine solutions (0.5 μL of 2, 4 6 8 and 10 mM) in buffer D (0.1 M sodium phosphate pH 8.0, 1 mM EDTA) were mixed with 9.5 μL buffer D and 10 μL DTNB (80 μg/mL) and incubated at room temperature for 15 mins. Three replicates of each condition were performed. Then the absorbance at 412 nm of each solution was measured in a UV spectrometer to generate a standard curve. Cyclic peptide TEV13 was dissolved in DMSO to achieve a stock concentration of 10 mM. The stock solution (0.5 μL) was mixed with 9.5 μL buffer D and 10 μL DTNB (80 μg/mL) and incubated at room temperature for 15 mins. Then the absorbance at 412 nm was measured in a UV spectrometer and fit to the standard curve generated with free cysteine.

### Competition of TEV13 for labeling by Cy5-TEV13

TEV13 (0.64 μL of 2 mM stock in DMSO) was added to 40 μL of TEV buffer and then 20 μL was taken to perform another 6 series of 2X dilutions. The 8^th^ condition is TEV buffer for use as the zero standard. Then 20 μL of 400 nM TEV protease and 40 μL of cell lysates were added to each well and the samples were incubated at 30 °C for 1 hour. Then 0.8 μL of 0.1 mM Cy5-TEV13 fluorescent probe was added to each condition and incubated at 30 °C for another hour before 30 μL 4X SDS PAGE sample buffer was added to each well to quench the reaction. After boiling the samples for 5 mins, 15 μL of the samples from each condition was analyzed on a 12 % SDS PAGE gel. Cy5 fluorescence was measured by scanning of the gel with a Typhoon 9410 variable mode imager (GE Healthcare, λ_ex_=633 nm, 670 BP30 filter). The fluorescent intensity of the protein bands at the expected size was analyzed using LI-COR Image Studio Lite software, and plotted with OriginPro 2019 software (OriginLab Corporation). Afterwards, the gel was stained with Coomassie blue to visualize the total protein load.

### Plasma stability of TEV13

TEV13 (10 μL of 2 mM stock in DMSO) was added to 290 μl mouse serum (final peptide concentration was 66.7 μM in 300 μL final volume in Balb C Mouse Serum (Innovative Research)) and the mixture was incubated at 37 °C in a water bath. After 0, 1, 2, 4 and 12 hours, 50 μL samples were taken, mixed with 100 μL analytical ethanol and kept at −20 °C for at least 1 hour to precipitate the proteins and lipids in plasma. Then the precipitants were centrifuged in an Eppendorf 5417 centrifuge at 14k rpm at 4 °C and the supernatant containing the peptides was mixed with 300 μL water before being lyophilized to dryness. Finally, the samples were resuspended in 55 μL water and 15 μL was characterized by analytical reversed-phase high performance liquid chromatography (RP-HPLC) on an Agilent HPLC system (1100 series) equipped with a Waters Agilent ZORBAX 300SB-C3 reverse phage column. The elution linear gradient was 0 - 95% v/v solvent B over 14 minutes at a flow rate of 1 mL/min (A: 99.9% v/v H2O and 0.1% v/v TFA; B: 99.9% v/v ACN and 0.1% v/v TFA). The elutes were detected at a wavelength of 220 nm.

### HPLC analysis of peptide purity

Peptide stock solutions (1 μL of 3 mM) were injected into an Agilent 1260 Infinity HPLC system equipped with a Poroshell 120 EC-C18 2.7 μm (Agilent #699975-902) analytical column, and run with a linear gradient of a mobile phase composed of eluent A (99.9 % v/v H2O, 0.1% v/v TFA) and eluent B (99.9% v/v acetonitrile, 0.1% v/v TFA) from 5% to 95% over 6 minutes at a flow rate of 0.6 ml/min. The absorbance at a wavelength of 220 nm was used to generate plots of peptide purity.

### Amber MD simulation

Homology models of TEV protease with TEV13 residues Glu6, Pro7, Leu8 and Tyr9 bound in the standard mechanism to the S6, S5, S4 and S3 sub-sites were built using X-ray cocrystal structure data sets using 1LVM (available at protein data bank) as a template. Non-essential water molecules and ions were deleted in the PDB files and the peptide ligands truncated to retain only the P6-P3 (ENLY) residues. The remaining residues in the peptide chain were added manually in randomly extended conformation. The initial structure of the DCA-VS linker was minimized with the Gaussian 16 software package and parameterized with GAFF. Tleap was used for generating molecular dynamic simulation input files, and the linkages between DCA-VS and cysteines residues in TEV13 were manually specified. Two individual models were built when the linkage between vinyl sulfone and active site cysteine was generated in R and S conformations. Force field parameters of DCA-VS and surrounding atoms were taken from GAFF and partial charges were calculated with HF/6-31G* RESP charge method. All structures were first minimized by holding the conformation of TEV13 Glu6, Pro7, Leu8 and Tyr9 residues and TEV protease, followed by whole system minimization. Then the system was heated to 300 K before equilibration on the whole system. The production equilibration was run at 300 K with constant pressure and periodic boundary for 40 ns. Langevin dynamics was used to control the temperature using a collision frequency of 1.0 /ps. Then the final structures were analyzed, the lowest energy structure is presented. All the calculations were performed by Assisted Model Building with Energy Refinement (AMBER, code available at http://ambermd.org/) biomolecular simulation package compiled in the sherlock High-Performance Computing (HPC) cluster provide by Stanford University and the Stanford Research Computing Center.

### Preparation of RAW cell extracts and labeling with Cy5 probes

RAW 264.7 cells (ATCC® TIB-71™) were cultured in DMEM (Gibco, #11995-073) supplemented with 10% animal serum (GeneMate, #S-1200-500), 100 units/mL penicillin (Gibco) and 100 µg/mL streptomycin (Gibco). Cells were cultured in a 5% CO2 humidified incubator at 37°C on 10 cm culture dish (corning, # 430167). At around 90% confluency, cells were scraped off into 1 mL 100 mM NaAc pH 5.5, 2.5 mM EDTA, 2 mM DTT buffer or 1 mL PBS pH 7.4 buffer supplemented with 2 mM DTT followed by lysis by ultrasound sonication. After centrifugation at 18,000xg at 4 °C for 15 mins, the supernatant was taken and aliquoted into 60 μL fractions. The probes (0.6 μL) were added to the lysates to achieve final probe concentrations of 1 μM, 0.5 μM, 0.25 μM, and 0.125 μM and incubated at 37 °C for 1 hour. After adding 15 μL SDS-PAGE sample buffer and bioloing, 15 μL of each labeling mixtures was analyzed on a 12 % SDS PAGE gel. Cy5 fluorescence was measured by scanning of the gel on a Typhoon 9410 variable mode imager (GE Healthcare, λ_ex_=633 nm, 670 BP30 filter). The fluorescent intensity of the labeled proteins at expected size was analyzed using the LI-COR Image Studio Lite software, and plotted with OriginPro 2019 software (OriginLab Corporation). Afterwards, the gel was stained with Coomassie blue to visualize the total proteins.

### Cat S, Cal L and SENP1 inhibition

The inhibitory activity of peptides against cathepsin S (Cat S), cathepsin L (Cat L) and sentrin-specific protease 1 (SENP1) was determined by pre-incubating peptide inhibitors with proteases and quantifying their residual activity with a fluorogenic substrate in a same procedure as described for determining TEV inhibition activities. For Cat S and Cat L inhibition, a final concentration of 2-200 μM peptides were pre-incubated with 2.5 nM Cat S or Cat L enzyme at 37 °C for 1 hour before 100 μM Z-FR-AMC substrate was added in 100 mM NaAc, 2.5mM EDTA, 2.5mM DTT, 0.01% Tween20, 1% DMSO buffer. Monitoring the generation of free AMC due to hydrolysis of fluorogenic substrate was achieved by reading fluorescent signal excited at 380 nm and emission at 460 nm. For SENP1 inhibition, final concentrations of 2-200 μM peptides were pre-incubated with 500 nM SENP1 enzyme at 37 °C for 1 hour before 100 μM Ac-QTGG-AFC substrate was added in 50 mM Tris 7.5, 100 mM NaCl, 1 mM EDTA, 1 mM DTT, 5% DMSO, 0.01% Tween20 buffer. Monitoring the generation of free AMC due to hydrolysis of fluorogenic substrate was achieved by reading fluorescent signal excited at 400 nm and emission at 505 nm. The slope of the developing curve reflected the residue activity of enzymes after covalent inhibition. The IC_50_ values were calculated as described above.

### Probe concentration and incubation time dependent labeling

To four tubes of 200 μL 100 nM TEV protease in 50 mM Tris-Cl, pH 8.0, 100 mM KCl, 1 mM EDTA 1 mM DTT and 0.01% Tween X100 buffer was added 2 μL of 100 μM, 50 μM, 25 μM, 12.5 μM peptide inhibitors in DMSO to achieve finaol peptide inhibitor concentrations of 1 μM, 0.5 μM, 0.25 μM and 0.125 μM. The reaction tubes were immediately transferred to 30 °C incubator, and after 5, 10, 20, 30, 40, 50, 60 mins, 20 μL of the reaction mixture from each condition was mixed with 10 μL SDS-PAGE sample buffer and boiled. Samples from each condition (15 μL) were further resolved on a 12 % SDS PAGE gel. Cy5 fluorescence was measured by scanning of the gel on a Typhoon 9410 variable mode imager (GE Healthcare, λ_ex_=633 nm, 670 BP30 filter). The fluorescent intensity of the labeled proteins at the expected size was analyzed using the LI-COR Image Studio Lite software, and plotted with OriginPro 2019 software (OriginLab Corporation).

