## Supplemental Figures for "A phage display approach to identify highly selective covalent binders"

**Supplementary Figures**

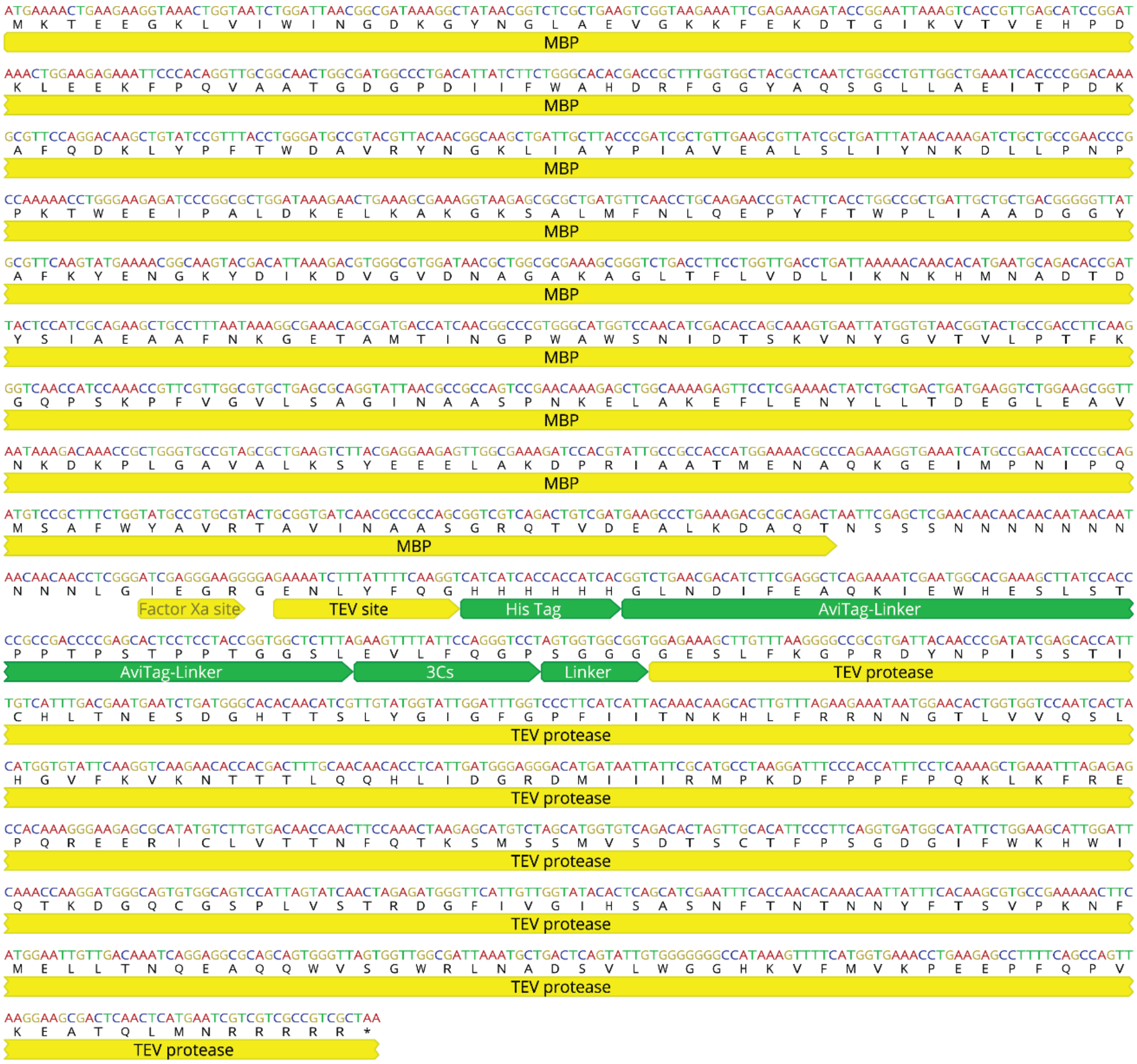


**Supplementary Fig. 1:** Expression construct for producing biotinylated TEV protease using the AviTag. The bacteria expression vector is built on the pMalC2 backbone. To improve the solubility of TEV protease in the cytosol of the expression host, an MBP domain is fused at the N-terminus of the expression construct. A TEV protease auto-cleavable peptide sequence (ENLYFQ|G) is placed before the expected TEV protease to allow the removal of MBP after translation. The desired TEV protease domain is N-terminally conjugated with a HisTag (for affinity purification), a Biotin AviTag (for immobilizing TEV protease on solid support), a rigid proline linker (to improve BirA biotinylation efficiency), a 3C protease cutting sequence (to allow orthogonally release TEV protease from solid support with 3C protease) and a flexible linker (to achieve efficient release TEV protease from solid support with 3C protease). Data Availability at GenBank accession numbers 2264028.

**a**

| Name | Amino acid sequences | IC50 (μM) |
|---|---|---|
| TEV16 | A C F V L E P L Y I C G | 2.57 ± 1.00 |
| TEV17 | A C Y V L E P L Y I C G | 4.69 ± 0.99 |
| TEV18 | A C H V L E P L Y I C G | 4.98 ± 1.18 |
| TEV19 | A C I V L E P L Y I C G | 4.56 ± 0.61 |
| TEV20 | A C D V L E P L Y I C G | 4.69 ± 0.54 |
| TEV21 | A C N V L E P L Y I C G | 6.29 ± 2.40 |
| TEV22 | A C K V L E P L Y I C G | 2.73 ± 0.59 |
| TEV23 | A C S V L E P L Y I C G | 6.35 ± 3.37 |
| TEV24 | A C M V L E P L Y I C G | 5.18 ± 1.74 |

**b**

| Name | Amino acid sequences | IC50 (μM) |
|---|---|---|
| TEV25 | A C W L L E P L Y I C G | 7.29 ± 1.13 |
| TEV26 | A C W I L E P L Y I C G | 5.36 ± 0.25 |
| TEV27 | A C W F L E P L Y I C G | 6.37 ± 0.56 |
| TEV28 | A C W Y L E P L Y I C G | 5.67 ± 0.61 |
| TEV29 | A C W W L E P L Y I C G | 7.94 ± 0.37 |
| TEV30 | A C W N L E P L Y I C G | 5.20 ± 0.67 |
| TEV31 | A C W D L E P L Y I C G | 4.72 ± 0.22 |
| TEV32 | A C W S L E P L Y I C G | 4.87 ± 0.06 |
| TEV33 | A C W M L E P L Y I C G | 1.53 ± 0.61 |
| TEV34 | A C W K L E P L Y I C G | 9.20 ± 1.12 |

**c**

| Name | Amino acid sequences | IC50 (μM) |
|---|---|---|
| TEV35 | A C W V V E P L Y I C G | 4.22 ± 0.86 |
| TEV36 | A C W V I E P L Y I C G | 11.21 ± 2.27 |
| TEV37 | A C W V F E P L Y I C G | 2.79 ± 0.31 |
| TEV38 | A C W V Y E P L Y I C G | 4.06 ± 2.53 |
| TEV39 | A C W V W E P L Y I C G | 5.11 ± 1.24 |
| TEV40 | A C W V Q E P L Y I C G | 2.5 ± 0.79 |
| TEV41 | A C W V E E P L Y I C G | 4.19 ± 0.48 |
| TEV42 | A C W V S E P L Y I C G | 4.09 ± 0.71 |
| TEV43 | A C W V M E P L Y I C G | 7.33 ± 2.62 |
| TEV44 | A C W V K E P L Y I C G | 9.61 ± 1.26 |

**d**

| Name | Amino acid sequences | IC50 (μM) |
|---|---|---|
| TEV45 | A C W V L E P L Y V C G | 4.63 ± 0.68 |
| TEV46 | A C W V L E P L Y F C G | 4.58 ± 1.30 |
| TEV47 | A C W V L E P L Y W C G | 4.65 ± 0.64 |
| TEV48 | A C W V L E P L Y Y C G | 8.82 ± 1.07 |
| TEV49 | A C W V L E P L Y H C G | 10.47 ± 1.53 |
| TEV50 | A C W V L E P L Y D C G | >100 |
| TEV51 | A C W V L E P L Y K C G | 10.45 ± 2.73 |
| TEV52 | A C W V L E P L Y R C G | 9.59 ± 1.79 |
| TEV53 | A C W V L E P L Y T C G | 4.41 ± 0.37 |
| TEV54 | A C W V L E P L Y Q C G | >100 |

**Supplementary Fig. 2.** Sequence activity relationship (SAR) study of cyclic peptide TEV3 by mutating residues at position (a) 3, (b) 4, (c) 5 and (d) 10 with amino acids of different properties of hydrophobicity, charge and size. The potency of each mutant for TEV protease is listed to the right as an $IC_{50}$ value.

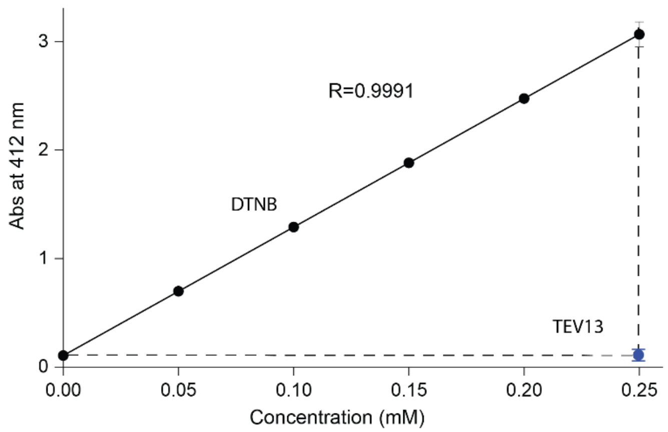


**Supplementary Fig. 3.** Ellmen’s test to determine the extent of unmodified cysteine on the cyclic peptide ABP TEV13. According the standard curve generated using free cysteine, no signification amount of free thiol is present in 0.25 mM cyclized peptide TEV13.

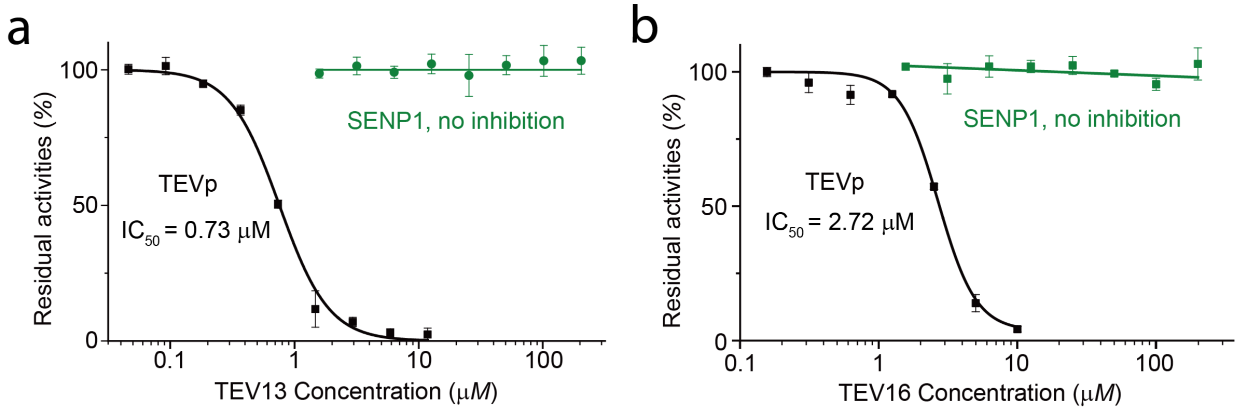


**Supplementary Fig. 4.** Cyclic peptide TEV13 and linear peptide TEV16 can inhibit TEV protease but not SENP1. Inhibition curves for TEV13 left and TEV16 right for TEV protease as well as recombinant SENP1.

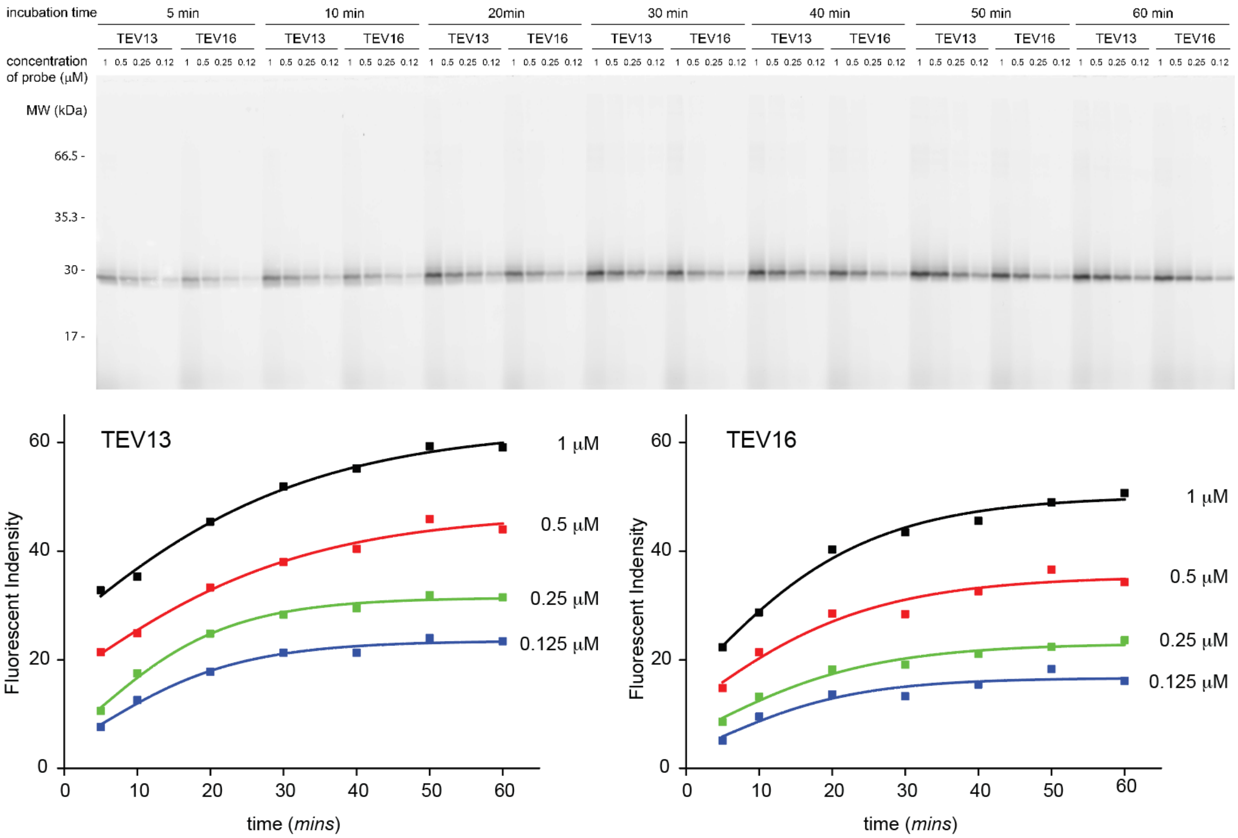


**Supplementary Fig. 5.** Labeling of TEV protease by fluorescently labeled TEV13 and TEV16 confirms that TEV3 is a faster and more effective label. TEV protease (100 nM) was incubated with increasing concentrations of the indicated fluorescent probes at 30 °C for the indicated times. After stopping the labeling reactions by adding SDS PAGE gel sample buffer and boiling, the labeling mixtures were analyzed by gel electrophoresis and detected by florescent scanning with a Typhone flatbed fluorescent scanner using the cy5 channel. The fluorescent intensity of cy5 labeled TEV protease was determined and plotted versus labeling time.

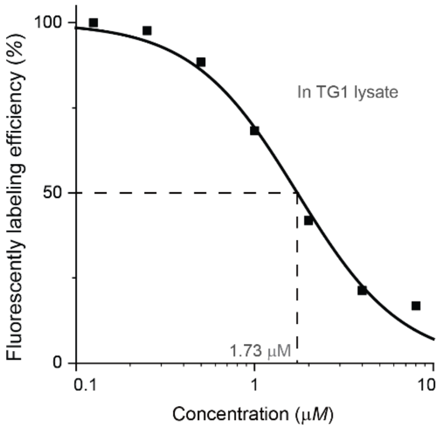


**Supplementary Fig. 6.** Competition of Cy5-TEV13 labeling of TEV protease by TEV13 in TG1 cell lysates. Plot shows values for the percent residual labeling by Cy5-TEV13 for the indicated TEV13 concentrations. The resulting $IC_{50}$ for inhibition of labeling is shown.

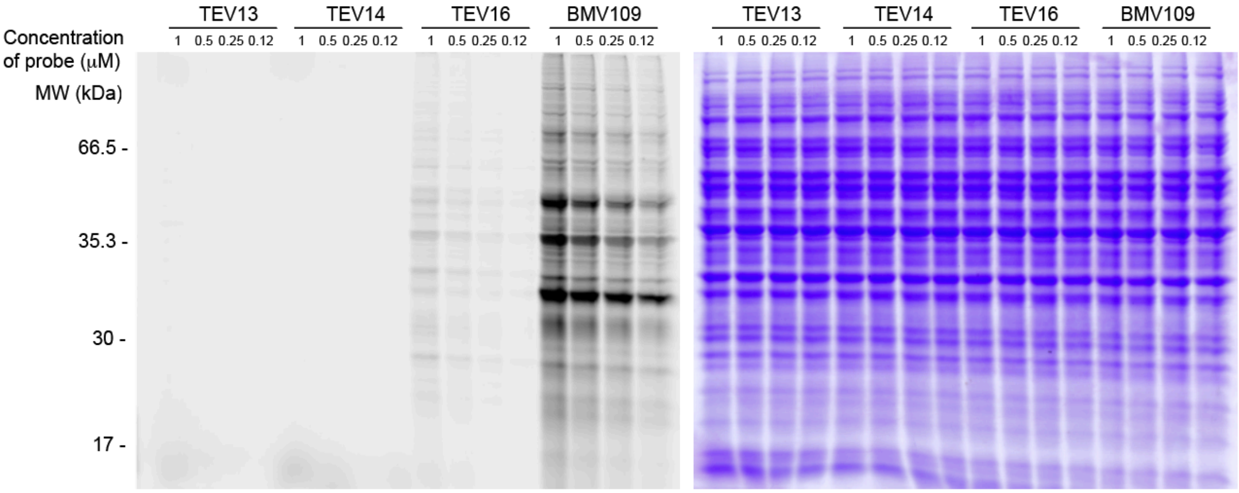


**Supplementary Fig. 7.** Fluorescently labeling of total protein extracts from RAW cells at pH7.4 with various probes. Images of SDS-PAGE gels containing samples of the indicated total cellular lysates labeled for 1 hr with the indicated Cy5-ABPs. Samples were separated on SDS-PAGE gels followed by scanning for Cy5 fluorescence using a flatbed laser scanner (Left) and then stained with Coomassie brilliant blue (CBB; right) to visualize total protein loading. The probe BMV-109 is a general ABP for cysteine cathepsins that was included for reference.

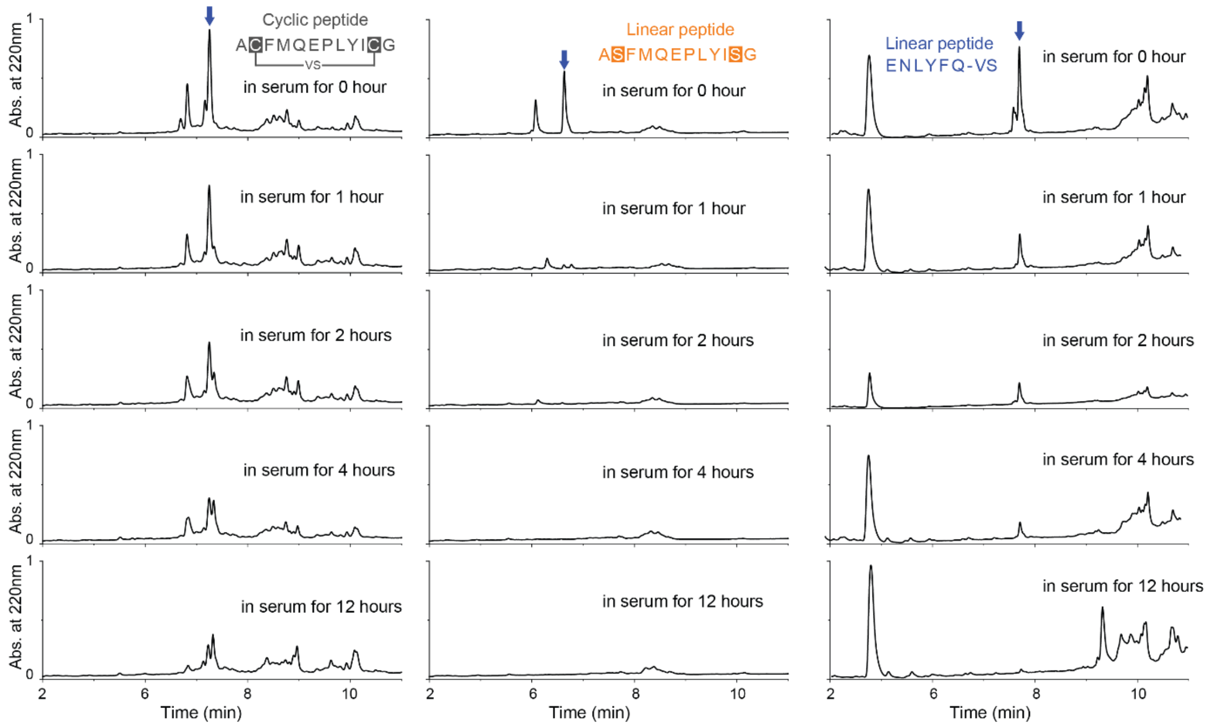


**Supplementary Fig. 8.** Plasma stability of cyclic peptide TEV13, a linear version of the TEV13 peptide (H-ASFMQEPLYISG-$NH_2$, serine residues were used to replace cysteine residues to avoid the oxidative formation of cyclic peptide with disulfide bridge) and the linear vinyl sulfone peptide TEV16. Plots are HPLC traces of the extracted peptides after incubation in mouse plasma at 37 °C for the indicated times. The location of the peak corresponding the intact original peptide is shown with a blue block arrow as determined by MS analysis.

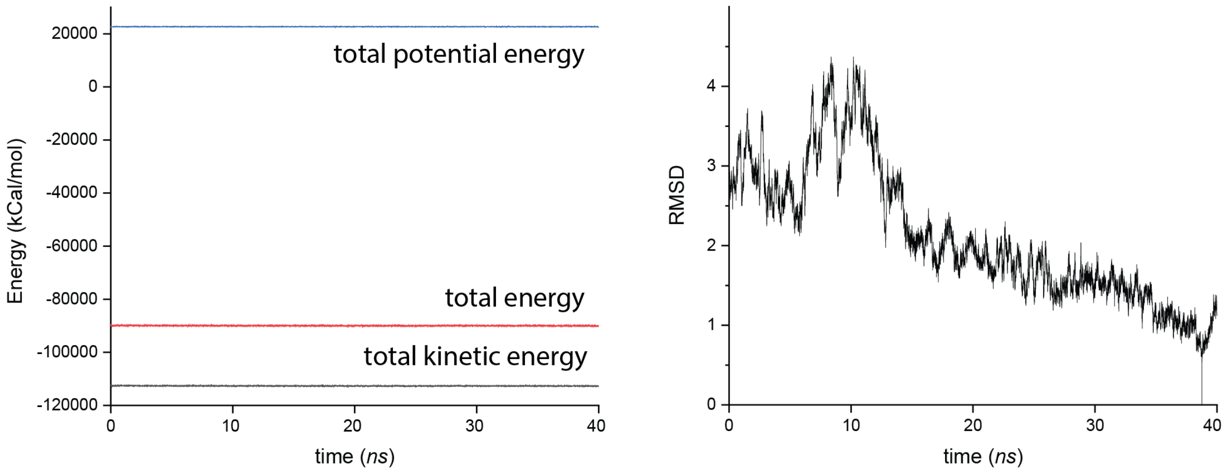


**Supplementary Fig. 9.** Analysis of Amber simulated structure of TEV13 complexed with the TEV protease. A) Total system energy of the 40 ns production simulation of the complex simulation in explicit TIP3P solvent models with ff14SB and gaff parameter sets. B) RMSD plot shows the evolution of the complex structure approaching the lowest energy conformation.

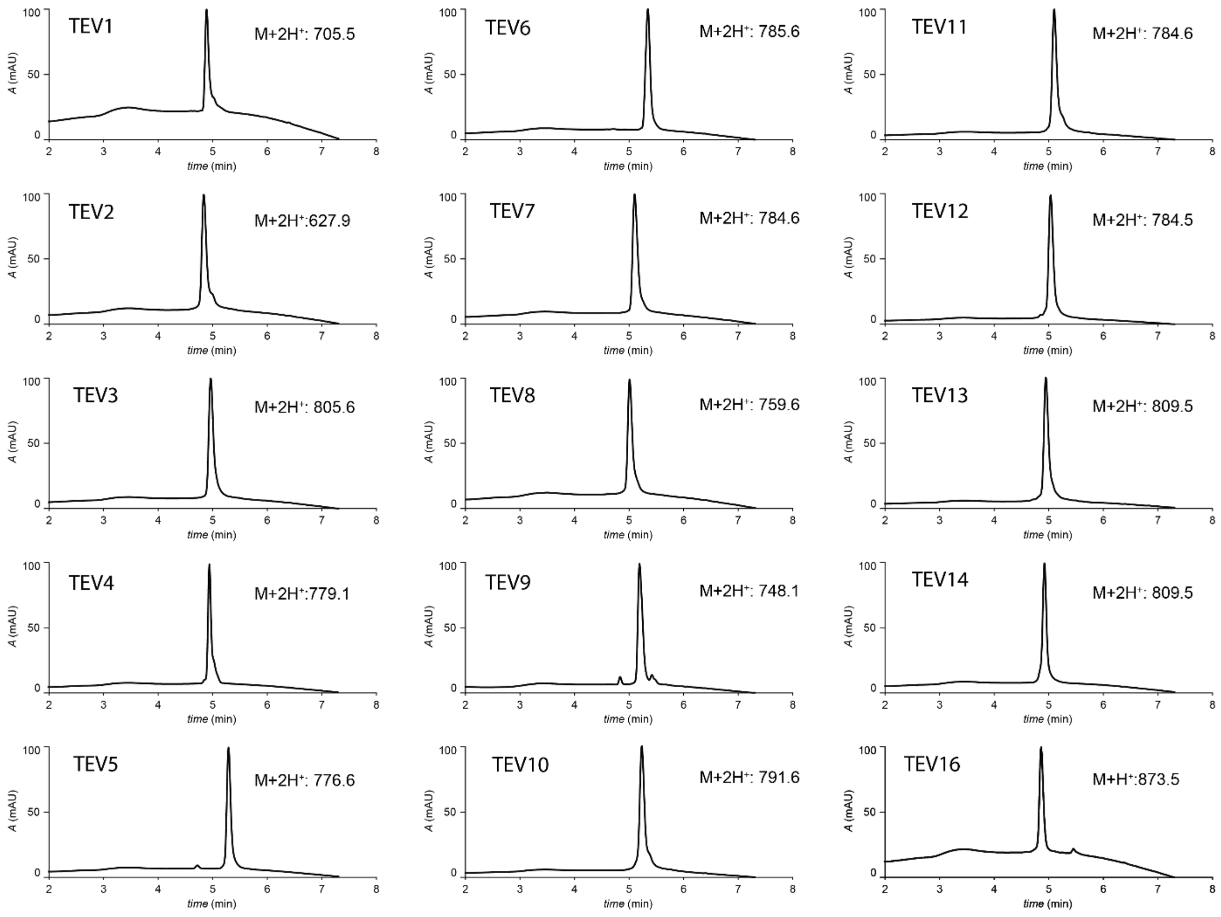


**Supplementary Fig. 10.** HPLC analysis of bicyclic peptides listed in Fig. 3. 1 μL of 3 mM peptide stock solutions are injected into an Agilent 1260 Infinity HPLC system for resolving, and spectrums of the absorbance at a wavelength of 220 nm and observed ESI MS of the corresponding peaks are presented.

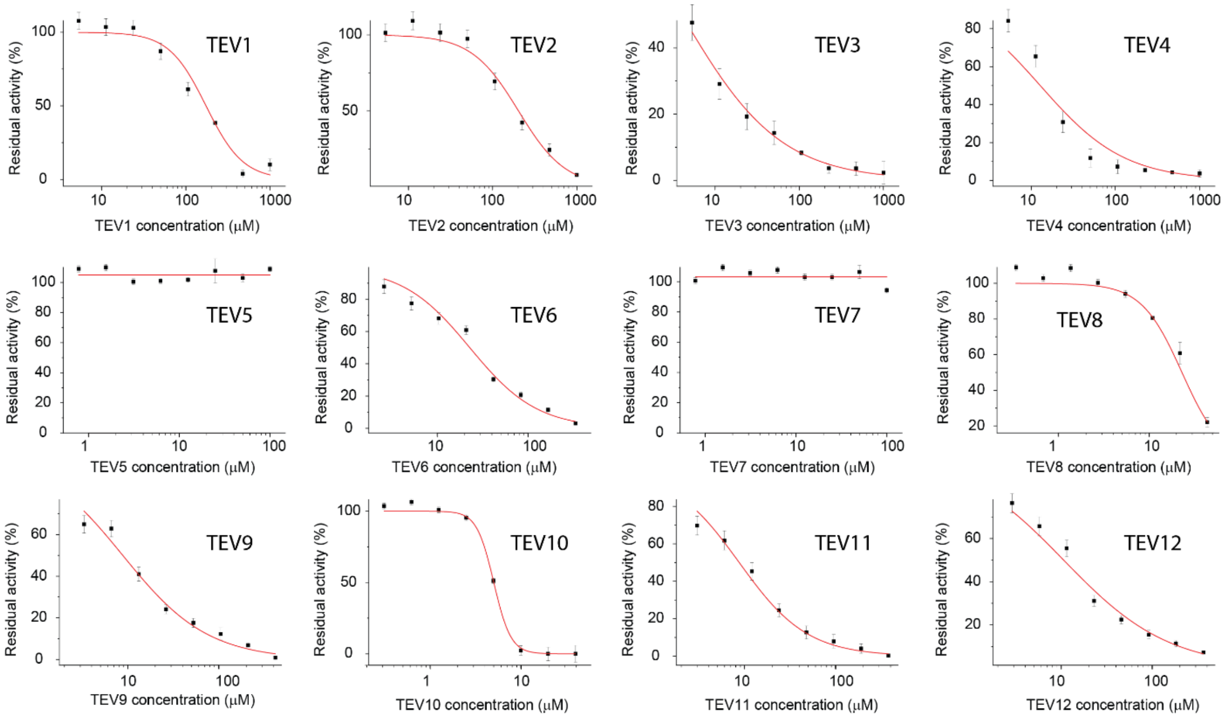


**Supplementary Fig. 11.** Dose response inhibition studies of the cyclized peptides TEV1 to TEV12 (Fig. 4) using the recombinant TEV. Plots show residual enzyme activity over a range of inhibitor doses as measured by hydrolysis of a fluorogenic substrate. Enzyme was pre-treated with inhibitor at the indicated concentrations for 1 hour followed by addition of substrate and measurement of activity. $IC_{50}$ values are indicated.
